# Diet-dependent sleep modulation by the *Drosophila* amino acid transporter ANIDRA

**DOI:** 10.64898/2026.05.20.726708

**Authors:** Ratna Chaturvedi, Rita R. Fagan, Chenghao Chen, Tobias Stork, Marc R. Freeman, Haley E. Melikian, Patrick Emery

## Abstract

Sleep is a conserved animal behavior necessary for survival. It is under tight circadian and homeostatic control, and modulated by diet. Here, we identify the amino acid transporter ANIDRA (ANID) as an important sleep regulator in *Drosophila*. Flies lacking ANID show decreased and poorly consolidated daytime and nighttime sleep. Contrary to wild-type controls, *anid* mutant flies are unable to adjust their sleep to their diet, behaving as if they were constantly on a complete diet rich in amino acids. ANID is expressed in ensheathing and cortex glia, where it inhibits mTOR activity in a diet-dependent manner. Moreover, pharmacological inhibition of mTOR attenuates the *anid* mutant sleep phenotypes. Interestingly, DH44-expressing brain neurons, which promote arousal and sense amino acids, are constantly active in ANID’s absence. We therefore propose that ANID mediates detection of dietary amino acids by ensheathing and cortex glia to regulate the activity of arousal-promoting neurons.

## Introduction

Among the most striking features of animal life is the universal need to sleep. Despite leaving organisms immobile and largely unresponsive to their environment, a state that carries substantial predation risk, sleep cannot be chronically curtailed without severe physiological and cognitive consequences. ^1–3^. Indeed, sleep supports a broad range of processes, from metabolic regulation and glymphatic clearance of waste products, to synaptic homeostasis, memory consolidation, tissue repair, and immune defense.

Sleep is controlled primarily by two interacting processes^4,5^. The circadian clock determines the timing of sleep (process C), while a homeostat determines the amount of sleep, based on previous wake activity (process S). The circuits by which the circadian clock regulates sleep are reasonably well known in both mice and *Drosophila*^3,6–8^. The homeostatic circuits appear to be quite complex and might involve multiple brain structures in both insect and mammals^3,7,9^. Sleep also responds to environmental inputs. For example, light pulses or mechanical stimuli can arouse animals^10,11^. Temperature impacts how much animals sleep during the day and night^12,13^. Internal states are important as well, stress and pain reduce sleep^14,15^, and the diet an animal is exposed to can profoundly impact sleep duration and architecture^16–19^.

Sleep and metabolism are deeply intertwined^20–22^. Insufficient sleep impairs insulin sensitivity and disrupts the hormonal regulation of appetite and energy balance, linking chronic sleep loss to metabolic dysfunction^21,22^. On the other hand, the brain’s nutritional state impact the activity of sleep circuits^23^. Among the dietary factors that influence sleep, amino acids have emerged as particularly important signals. Beyond their well- known role as biosynthetic precursors for key neurotransmitters (i.e. serotonin, dopamine, glutamate, and GABA) amino acids are increasingly recognized as direct nutrient signals impacting the activity of sleep circuits^23^.

*Drosophila* is a powerful model to understand the mechanism generating and controlling sleep, because of its genetics, refined tools to image and manipulate neural circuits, and the ease with which environmental conditions and internal states can be adjusted to understand their impact on sleep^2,3^. Multiple studies have probed the impact of diet on *Drosophila* sleep. Dietary sugars promote sleep as long as they stimulate sweet taste receptors^24,25^. On the other hand, a complete diet with high concentrations of amino acids promotes wakefulness^17^. Studies systematically testing the role of single amino acids revealed that L-threonine promotes sleep through GABA signaling inhibiting R2 neurons in the Ellipsoid Body (EB), a central sleep regulating center^26^. However, L-leucine and L-phenylalanine were found to reduce sleep duration. L-threonine decreases sleep latency, while L-glutamine and L-glutamate increase sleep latency.

More recently, the impact of dextrogyre (D) amino acids on sleep was also tested, with D-serine and to a lesser extend D-glutamine promoting sleep^27^. Moreover, a recent study indicated that the amino acid LAT1-like transporters JHL21 and MND, expressed in the blood brain barrier, are important for maintaining proper brain amino acid levels and thus promote sleep via GABAergic signaling on arousal-promoting circadian clock neurons^28^. Interestingly, dietary amino acids protect sleep from the effect of age by preserving its consolidation^29^. DH44+ neurons, which promote wakefulness and feeding, increase their activity in response to amino acids with the help of the amino acid transporter CG13248 ^30–32^. This might explain the overall sleep-reducing effect of a diet rich in amino acids. Interestingly, the impact of amino acids on sleep can be modulated by sugars and whether they are sweet-tasting or not^33^.

Here, we identify a novel and critical mechanism by which diet modulates sleep in *Drosophila*. We show that the amino acid transporter ANIDRA (CG7888), expressed in ensheathing and cortex glia, promote sleep and its consolidation. It does so by inhibiting the mTOR pathway when flies are exposed to a nutrient-poor diet. When ANIDRA is missing, DH44+ neurons are strongly activated, which helps explaining why flies sleep less, and are unable to modulate their sleep duration and consolidation as a function of the diet they are exposed to. These findings reveal a glial amino acid-sensing mechanism that translates nutritional status into neural circuit activity to govern sleep.

## Results

### ANIDRA promotes sleep and its consolidation

In an unpublished expression study, the mRNA encoding CG7888, a putative amino acid transporter, was found to be enriched in astrocytes (A. N. Fox, personal communication). This was confirmed in single cell (sc) RNAseq studies, which indicated broader glial *cg7888* mRNA expression in Ensheathing and Cortex glia^34,35^. To study CG7888’s function, we generated a null mutant allele using CRISPR/Cas9-mediated genome editing (Figure 1A). To verify that *cg7888^del^* mutants indeed do not produce CG7888, we used a custom-made antibody to CG7888. As expected, no protein could be detected by Western Blot in mutant head protein extracts (Figure 1B), while a protein with an apparent molecular weight of 37 kDa (slightly smaller than the predicted 44 kDa) was present when probing wild-type (*w^1118^*) head extracts. CG7888 was also undetectable by immunohistochemistry in mutant fly brains (Figure 1C, a weak diffuse signal remains visible in the mutant, which must be the result of non-specific antibody binding since it is present throughout the brain and does not appear to vary across brain regions).

**Figure 1.**
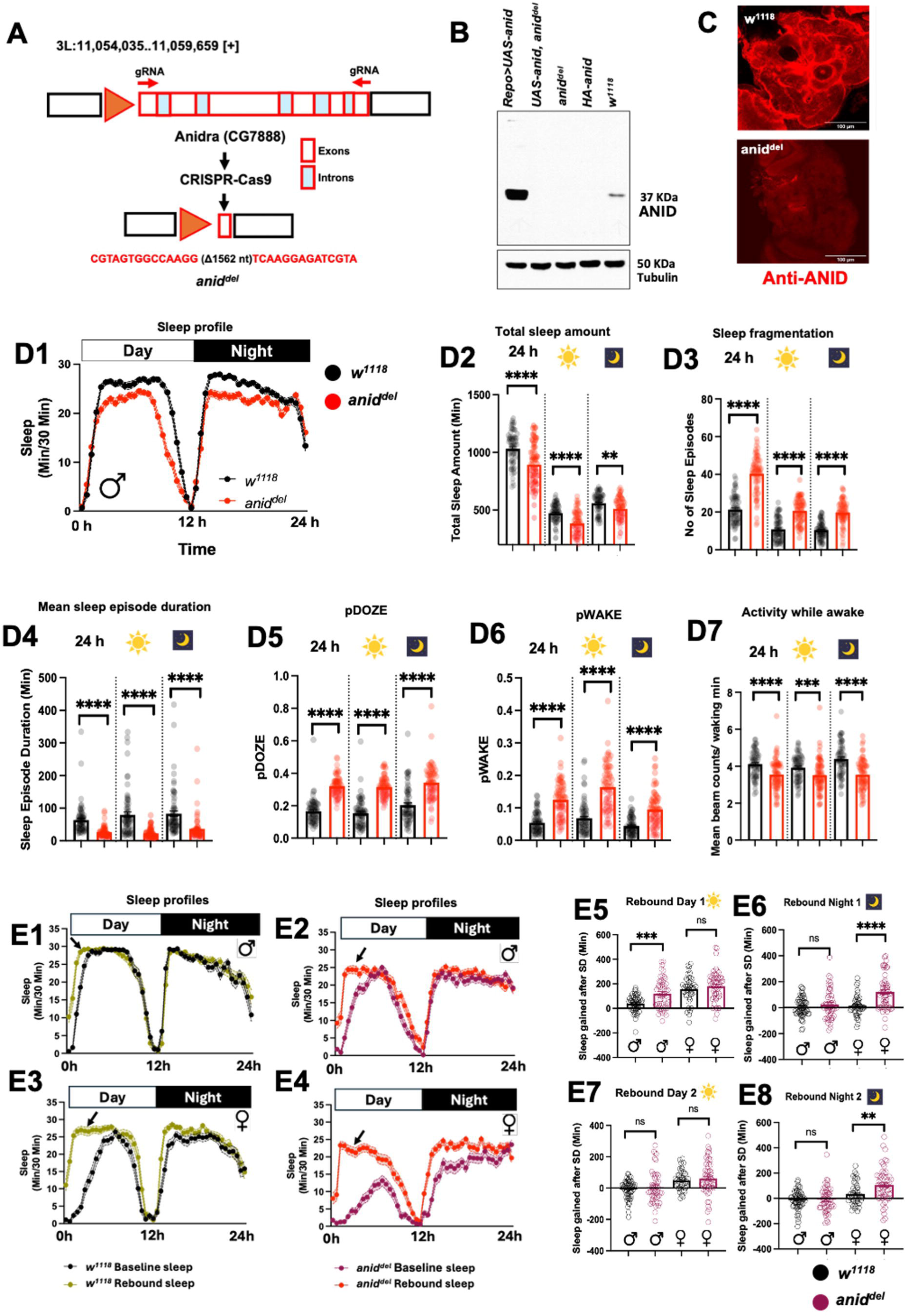
Loss of ANID disrupts sleep. **(A)** Schematic of CRISPR-mediated deletion strategy used to generate the *anidra* (*anid*, *cg7888*) null mutant (*anid^del^*). **(B)** Western blot validation of a rabbit polyclonal antibody raised against an ANID peptide (N-terminal). No ANID signal is detected in mutant flies, and signal is increased in flies overexpressing ANID. Signal is also very low in a HA-tagged allele that was generated by CRISPR/Cas9 genome editing. This allele was not used further in this study. **(C)** ANID signal is also lost in mutant flies when probed in adult *Drosophila* brain with immunohistochemistry. A weak, uniform background staining is visible in the mutant. **(D1- D7)** Sleep behavior in *w^1118^* (wild-type) and *anid^del^* male flies. (D1) Dailly sleep profiles of *anid^del^* (red; n = 69) and *w^1118^* (black; n = 68) males. (D2) Total sleep (min). (D3) Number of sleep episodes. (D4) Mean sleep episode duration (min). (D5) Probability of falling asleep (pDOZE; a measure of sleep pressure). (D6) Probability of waking (pWAKE; a measure of sleep maintenance). (D7) Mean locomotor activity expressed as beam breaks per waking minute. All parameters were quantified over 24 h and further subdivided into daytime and nighttime periods. **(E1- E8)** Enhanced sleep homeostasis in *anid^del^* mutants following sleep deprivation. Sleep profiles under baseline conditions and following 12-h of nighttime sleep deprivation (SD) for (E1) *w^1118^* males (n = 61), (E2) *anid^del^*males (n = 52), (E3) *w^1118^* females (n = 45), and (E4) *anid^del^* females (n = 56). Baseline is shown in black (*w^1118^*) or magenta (*anid^del^*); post-SD recovery is shown in olive green (*w^1118^*) or red (*anid^del^*). Black arrows in the sleep profile indicates extra sleep. (E5- E8) Sleep rebound quantified during day 1 (E5), night 1 (E6), day 2 (E7), and night 2 (E8) following SD. Data are presented as mean ± SEM. Statistical comparisons between *anid^del^* and *w^1118^* were performed using two-tailed Student’s t-test for normally distributed data or the Mann–Whitney U test for non-normally distributed data. ****p < 0.0001, ***p < 0.001, **p < 0.01, *p < 0.05; ns, not significant. See also Figure S1

*cg7888^del^* animals appeared healthy, their movements well-coordinated, and they were able to reproduce when grown on regular complete diet. Since glial cells contain circadian clocks and are implicated in the control of sleep in *Drosophila*^34,36–42^, we tested whether CG7888 impacts circadian behavior and sleep. Under a 12 h:12 h light/dark (L:D) cycle, *cg7888^del^*flies exhibited the typical circadian morning and evening anticipatory behavior of the lights-on and lights-off transitions, controlled by the M- and E-oscillators^43,44^ (Figure S1A1-2). Under constant darkness (DD), the period of circadian rhythms was close to 24h, though slightly elongated compared to wild-type controls (Figure S1A4). We therefore conclude that CG7888 has little impact on the circadian neural network. However, we did notice an increase in the fraction of arrhythmic flies (Figure S1A3), which could indicate a defect in neural pathways downstream of the circadian neural network (see below).

The daily sleep profile was clearly altered in mutant flies (Figure 1D). Indeed, total sleep duration was reduced both at night and during the day in *cg7888^del^* mutant flies (Figure 1D1-D2). Moreover, sleep was excessively fragmented. The number of sleep bouts were increased (Figure 1D3), their average duration was decreased (Figure 1D4), and the probability for flies to fall asleep (pDOZE) or wake up (pWAKE) was clearly increased (Figure 1D5-6). These phenotypes were not the result of general hyperactivity. In fact, mutant flies were slightly hypoactive when awake (Figure 1D7). All these phenotypes, observed in both males (Figure 1D) and females (Figure S1B), are indicative of reduced and poor sleep quality, and increased sleep pressure. We therefore propose to name CG7888 as ANIDRA (ANID), a Sanskrit word that translates to sleeplessness.

Since *anid^del^* mutants show such severely defective baseline sleep phenotypes, we wondered whether their homeostatic response to sleep deprivation (sleep rebound) is also disrupted. We therefore mechanically sleep-deprived control and mutant flies for 12 h during the night. Both mutant males and females showed greater sleep rebound than control flies, probably reflecting their chronic sleep deprivation (Figure 1E and Figure S1C). We noticed that in mutant females, the increased magnitude of sleep rebound resulted from elevated nighttime sleep rebound (Figure 1E6, E8), while in males daytime sleep rebound was increased (Figure 1E5). Thus, ANID’s impact on sleep homeostasis is sexually dimorphic. However, this study is focused on basal sleep, which is similarly affected in males and females. For the rest of the study, we will thus focus on male flies.

### ANIDRA functions in ensheathing and cortex glia

To begin to understand how ANID regulates sleep, we first determined its pattern of expression in the fly brain, using our ANID antibody and glial GAL4 drivers to mark different types of glia. ANID expression was predominantly glial, with low or no expression in neuropils (Figure 1C, S3D). Using GFP markers driven by specific GAL4 lines, we found ANID to be strongly expressed in Ensheathing glia and Cortex glia (Figure 2A, S2A). To our surprise ANID expression was not obviously detectable in astrocytes, even though the *anid* mRNA is present in these cells^34,35^ (Figure 2A). *anid* mRNAs might thus be inefficiently translated in astrocytes, or translated there only under specific environmental conditions. We did not observe obvious expression in surface glia (Figure 1C, S3D), in agreement with scRNAseq data^34^.

**Figure 2.**
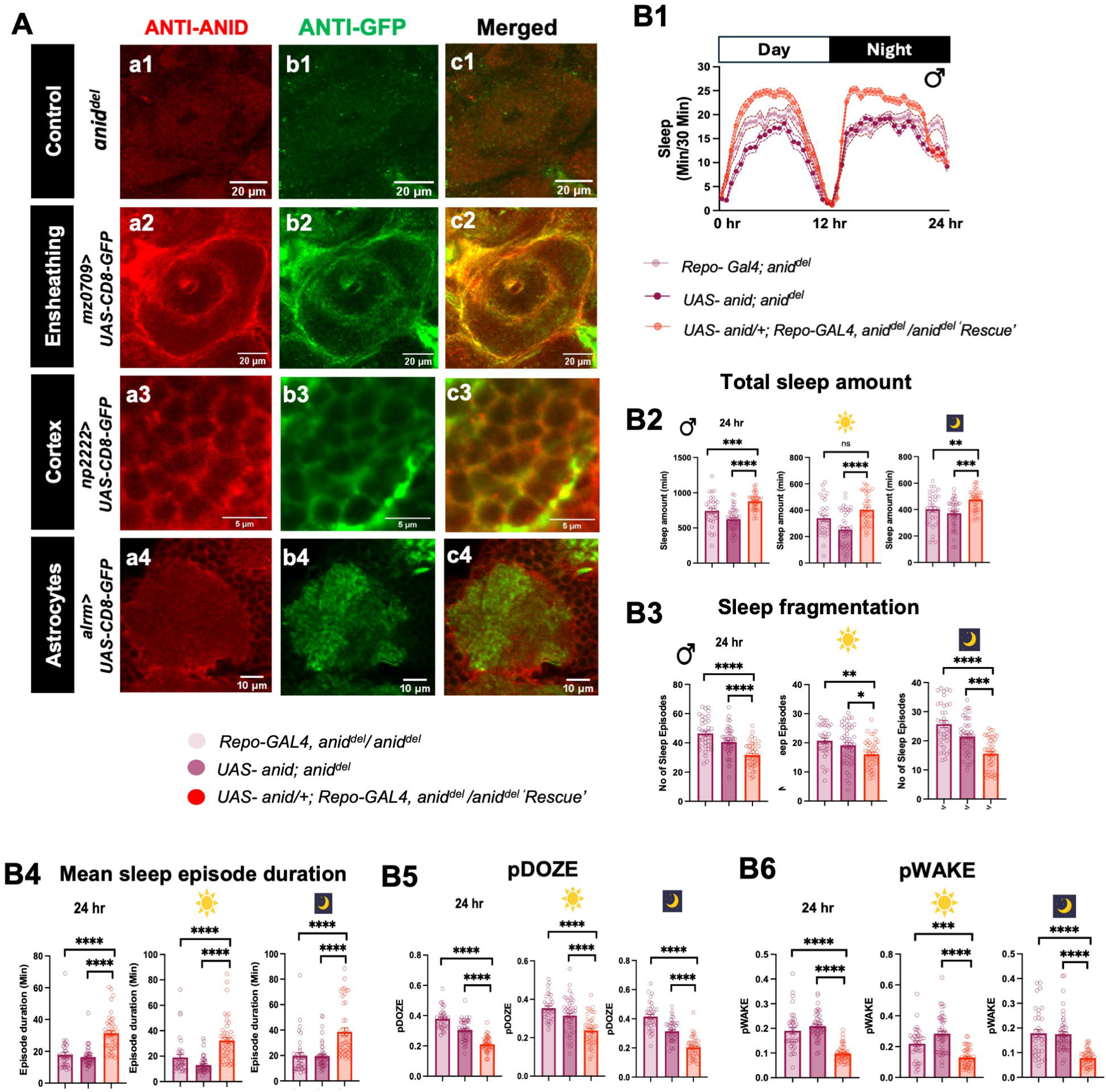
Glial ANIDRA expression controls sleep. **(A)** ANID and GFP immunostaining in adult *Drosophila* brains. CD8-GFP was expressed in distinct glial subtypes using specific GAL4 drivers. *anid^del^* mutant brains served as negative control. Columns a, b, and c represent anti-ANID (red), anti-GFP (green) and merged channels, respectively. (a1- c1) *anid^del^*, central complex region (a2- c2) Ensheathing glia labeled with *mz0709*-*GAL4* in the central complex region. (a3- c3) Cortex glia labeled with *np2222*-*GAL4*. (a4- c4) Astrocytic glia labeled with *Alrm*-*GAL4*. Scale bars are indicated in each panel. **(B1- B6)** Pan-glial expression of *anid* rescues sleep defects in *anid^del^* mutants. (B1) Daily sleep profiles of *UAS-anid*; *anid^del^* (magenta; n = 47), *repo*-*GAL4*, *anid^del^*(light magenta; n = 36), and *repo*-*GAL4* > *UAS-anid* in the *anid^del^* background (red; n = 45) male flies. (B2) Total sleep (min). (B3) Number of sleep episodes. (B4) Mean sleep episode duration (min). (B5) pDOZE. (B6) pWAKE. All parameters were analyzed over 24 h and during daytime and nighttime periods separately. Data are presented as mean ± SEM. Comparisons among genotypes were performed using ordinary one-way ANOVA for normally distributed data or a Kruskal- Wallis test for non-normally distributed data. ****p < 0.0001, ***p < 0.001, **p < 0.01, *p < 0.05; ns, not significant. See also Figure S2

To confirm that ANID functions in glia to regulate sleep, we expressed ANID in all glial subtypes of *anid^del^* flies using the pan-glial driver *repo-GAL4*. As expected, sleep amount and sleep quality were significantly improved compared to both mutant controls (Figure 2B, S2B), demonstrating that *anid* deletion was causative of the sleep phenotypes, and that ANID functions in glia to control sleep. Because ANID is also highly expressed in the hindgut according to Flybase, we dissected and stained guts with the ANID antibody. Indeed, we found ANID expression in hindgut enterocytes (Figure S2C), but *repo-GAL4* is not active in this tissue. Hence, the rescue of sleep with this driver is not the result of restoring ANID expression in the gut.

We observed that ectopic ANID expression could cause lethality, complicating spatial mapping of ANID function through a rescue approach. Thus, we turned to tissue-specific CRISPR/Cas9 genome editing to identify the glial subtype(s) in which ANID is (are) required to control sleep. We expressed two guide RNAs targeting *anid* with *repo-GAL4* and observed severe reduction of ANID expression in the central brain (Figure S3D), Pan-glial *anid* disruption recapitulated the *anid^del^* phenotypes: sleep duration was decreased, sleep fragmentation as well as pDOZE and pWAKE were increased, both at night and during the day (Figure 3A, S3A). Interestingly, ablation of *anid* in either ensheathing (Figure 3B, S3B) or cortex (Figure 3C, S3C) glia reduced sleep quality and total sleep amount. However, these phenotypes were milder than those observed when all glia were targeted. We thus conclude that ANID functions in both cortex and ensheathing glia to impact sleep, which fits with its strong expression in these two cell types. We cannot entirely exclude a minor contribution from other cell types where ANID might expressed at low levels, such as astrocytes.

**Figure 3.**
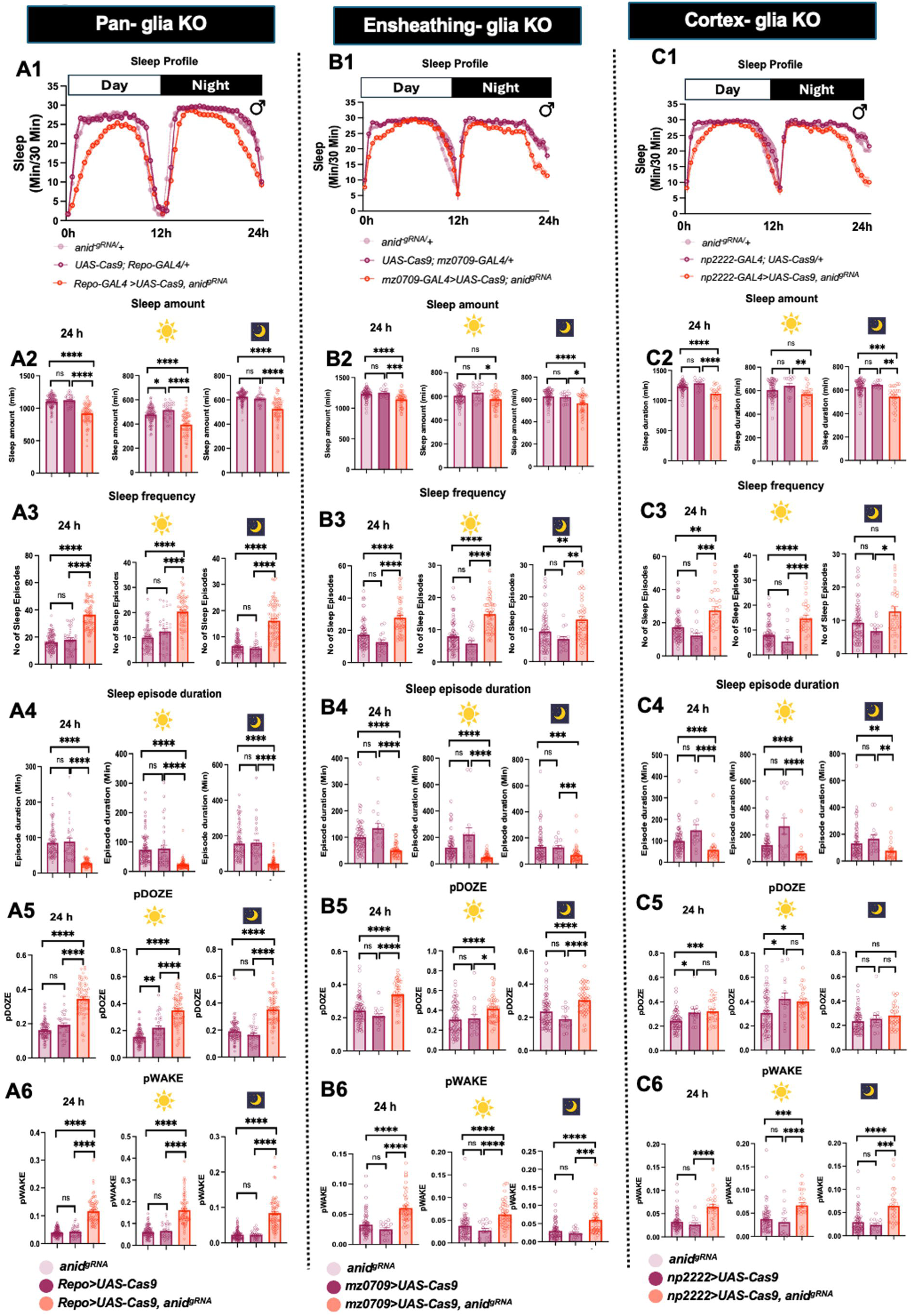
ANIDRA expression in both ensheathing and cortex glia contributes to sleep regulation. **(A1- A6)** Sleep defects following CRISPR-mediated deletion of *anid* in all glia using *repo*-*GAL4*. (A1) Sleep profiles of *UAS-anid*^gRNA^ (light magenta; n = 115), *UAS-Cas9*; *repo*-G*AL4* (magenta; n = 35), and *repo*-*GAL4* > *UAS-Cas9*; *UAS-anid^gRNA^* (red; n = 77) male flies. (A2) Total sleep (min). (A3) Number of sleep episodes. (A4) Mean sleep episode duration (min). (A5) pDOZE. (A6) pWAKE. **(B1- B6)** Sleep defects following CRISPR-mediated deletion of *anid* in ensheathing glia using *mz0709*-*GAL4*. (B1) Sleep profiles of *UAS-anid*^gRNA^ (light magenta; n = 77), *UAS-Cas9*; *mz0709*-*GAL4* (magenta; n = 20), and *mz0709*-*GAL4* > *UAS-Cas9*; *UAS-anid^gRNA^*(red; n = 48) male flies. (B2) Total sleep (min). (B3) Number of sleep episodes. (B4) Mean sleep episode duration (min). (B5) pDOZE. (B6) pWAKE. **(C1- C6)** Sleep defects following CRISPR-mediated deletion of *anid* in cortex glia using *np2222*-*GAL4*. (C1) Sleep profiles of *UAS-anid^gRNA^* (light magenta; n = 77), *UAS-Cas9*; *np2222*-*GAL4* (magenta; n = 20), and *np2222*-*G*AL*4* > *UAS-Cas9*; *UAS-anid^gRNA^*(red; n = 27) male flies. (C2) Total sleep (min). (C3) Number of sleep episodes. (C4) Mean sleep episode duration (min). (C5) pDOZE. (C6) pWAKE. All parameters were analyzed over 24 h and during daytime and nighttime periods separately. Data are presented as mean ± SEM. Statistics as described in Figure 2. See also Figure S3

### ANIDRA is an amino acid transporter

ANID is homologous to the proton-coupled SLC36 amino acid transporter family (PAT). Its close homolog in *Aedes aegypti* mosquitoes (AaePAT1) was found to be a low affinity and low specificity transporter for several amino acids^45^. We therefore stably expressed ANID in HEK293T cells and tested whether it indeed functions as a H^+^-dependent amino acid transporter. ANID expressed in mammalian cells was detected at the plasma membrane (Figure S4A), and also robustly in RAB11^+^ endosomes (Figure S4B). We first characterized ANID’s transport potential using radiolabeled proline (Figure 4A) and found that ANID exhibits saturable proline transport, with a significantly higher proline affinity at acidic vs. neutral pH (Figure 4A-C), confirming that ANID is a H^+^-dependent transporter. The mammalian SLC36 ortholog, PAT1 is competitively inhibited by 5-hydroxy-L-tryptophan (5HTP)^46^, and we found that 5HTP dose-dependently inhibited proline transport by ANIDRA with an IC_50_=2mM (Figure S4C), which is similar to its potency at PAT1. Kinetics studies revealed that ANID also exhibits saturable L-glutamate, L-alanine, and L-glutamine transport, and that sodium differentially impacts substrate transport, increasing proline affinity, but decreasing and/or severely compromising glutamate, alanine, and glutamine transport affinity. (Figure 4D-G, S4C-F and Table I). A single dose competition screen using unlabeled amino acids further revealed that multiple amino acids block proline transport via ANID, suggesting that ANID can, at minimum, bind these amino acids and that they may also be substrates (Figure S4C).

**Table 1.** ANIDRA uptake kinetics. K_m_ and V_max_ values ±S.E.M. as determined by [^3^H] substrate uptake in the presence or absence of Na+, pH 6.5 Values are presented in mM. N.F. = No fit

| [$^3$H]substrate | +$Na^+$ | | -$Na^+$ | |
| --- | --- | --- | --- | --- |
| | $V_{max}$<br>(pmol/min/mg) | $K_m$<br>(μM) | $V_{max}$<br>(pmol/min/mg) | $K_m$<br>(μM) |
| Proline | 6828 ± 2527 | 0.36 ± 0.04<sup>a</sup> | 7378 ± 1082 | 1.75 ± 0.14 |
| Alanine | N.F. | N.F. | 5835 ± 540 | 2.84 ± 0.51 |
| Glutamate | 20388 ± 4973<sup>b</sup> | 10.9 ± 3.0<sup>c</sup> | 3840 ± 1061 | 1.29 ± 0.31 |
| Glutamine | N.F. | N.F. | 8980 ± 2472 | 0.45 ± 0.24 |

<sup>a</sup>Significantly less than (-) $Na^+$ , $p < 0.0001$ , two-tailed, unpaired Students t test, N= 3 (- $Na^+$ ) and 9 (+ $Na^+$ ).
<sup>b,c</sup> Significantly greater than (-) $Na^+$ , $p < 0.01$ , two-tailed, unpaired Students t test, N= 4 (- $Na^+$ ) and 2 (+ $Na^+$ ).

**Figure 4.**
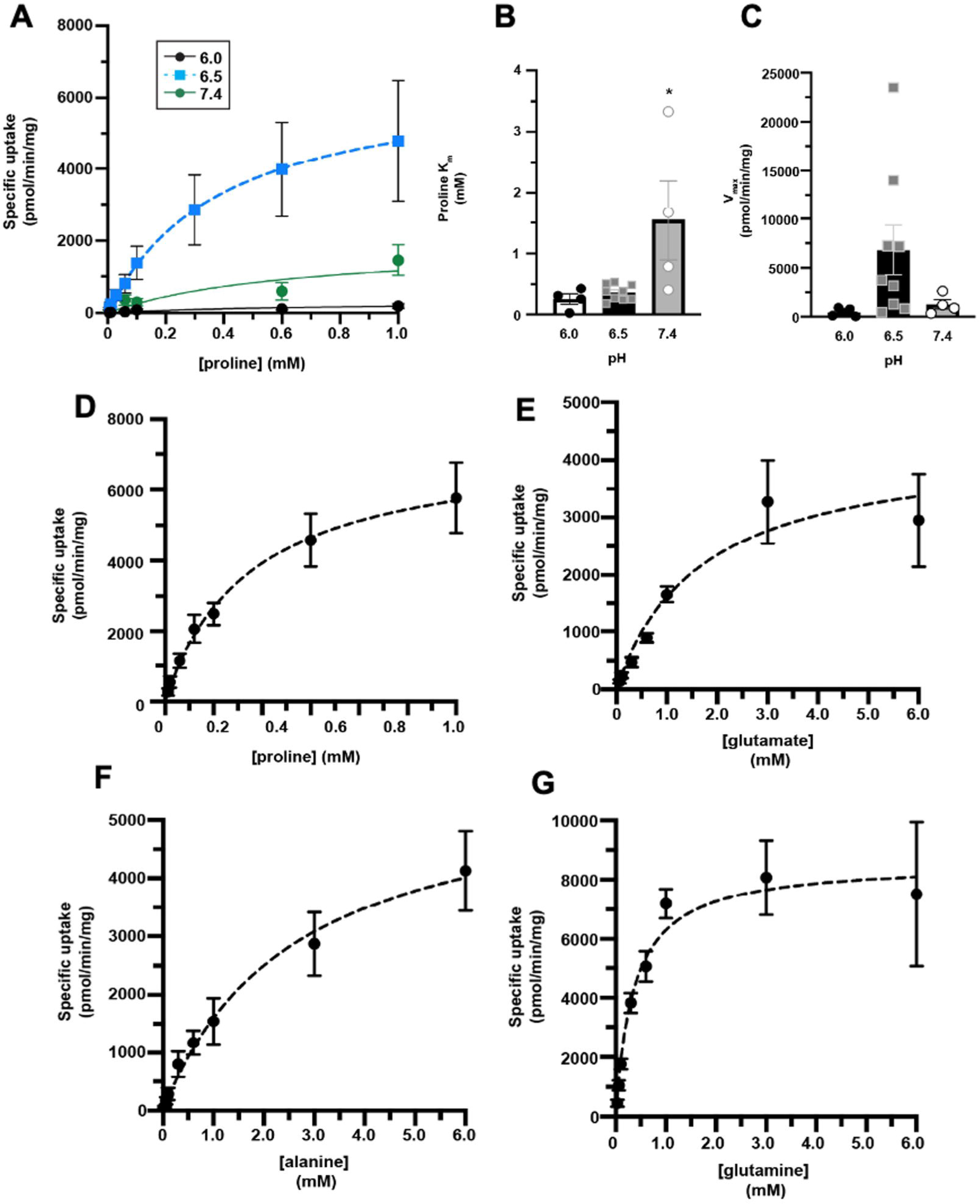
ANIDRA is a proton-activated amino acid transporter. *[^3^H]substrate uptake.* **(A- C)** Proline uptake by ANIDRA is saturable and pH-sensitive. Proline uptake assay in stably- transfected HA- ANIDRA HEK293T cells. [^3^H]proline uptake kinetics were measured at the indicated [proline] and pHs, and kinetic constants were calculated as described in Methods. **(A)** Saturation uptake at pH 6.0, 6.5, and 7.4. **(B)** ANIDRA substrate affinity for proline is pH-sensitive. *Significantly different from pH 6.5, p=0.01, one-way ANOVA test with Dunnett’s multiple comparison test. **(C)** ANIDRA proline V_max_ is pH_sensitive. Brown-Forsythe and Welch ANOVA test (p=0.03). N=4-9. **(D-G)** Saturation uptake kinetics for proline (D), glutamate (E), alanine (F) and glutamine (G), measured at pH 6.5, in the absence of Na^+^. K_m_ and V_max_ kinetic constants ±Na^+^ are reported in Table I.

### ANIDRA is required for diet-dependent sleep modulation

Since ANID transports amino acids, we wondered whether it modulates sleep in a diet-dependent manner. Our standard sleep assay is performed with a rather nutrient-poor food, containing only sucrose as a source of energy. We thus measured sleep when flies were exposed to a complete diet rich in amino acids (see methods). As previously reported^17^, we found that sleep duration was reduced in wild-type flies exposed to a complete diet, compared to sucrose-only diet. However, under our experimental conditions, this reduction was only observed at night. Sleep was also significantly deconsolidated during the night, with increased sleep bout numbers and decreased sleep bout length. A similar trend was observed during the day, but significance was not reached for either parameter. Strikingly, *anid^del^* flies did not modulate sleep as a function of diet, appearing to behave as if constantly on nutrient-rich food (Figure 5A-D). We wondered whether the decrease in sleep could be the result of *anid* mutant flies feeding excessively. However, when performing a CAFÉ assay^47^, we did not observe a change in food consumption when flies were of the same age and exposed for the same amount of time to sucrose food as flies tested for sleep (Figure S5D, S5E). However, we observed that *anid^del^* flies had a shorter lifespan on sucrose diet compared to complete diet, and compared to wild-type controls on sucrose-only food. They started to die after about 10 days when fed only sucrose (Figure S5A,B), spending most of their time on food as their abdomen became swollen (Figure S5F). Note that we measure sleep 4-6 days into exposure to sucrose food, days before flies start to die and at a time when *anid^del^* mutants consume as mentioned above a normal amount of food (Figure S5D-E), and that flies are healthy on nutrient-rich food but still show sleep phenotypes (Figure 5, S5B). Moreover, glial rescue of ANID expression only weakly rescued mortality on sucrose food (Figure 5SC), while it robustly rescued the sleep phenotypes (Fig. 2). Thus, the various sleep phenotypes we observed are not causally related to starvation or poor health.

**Figure 5.**
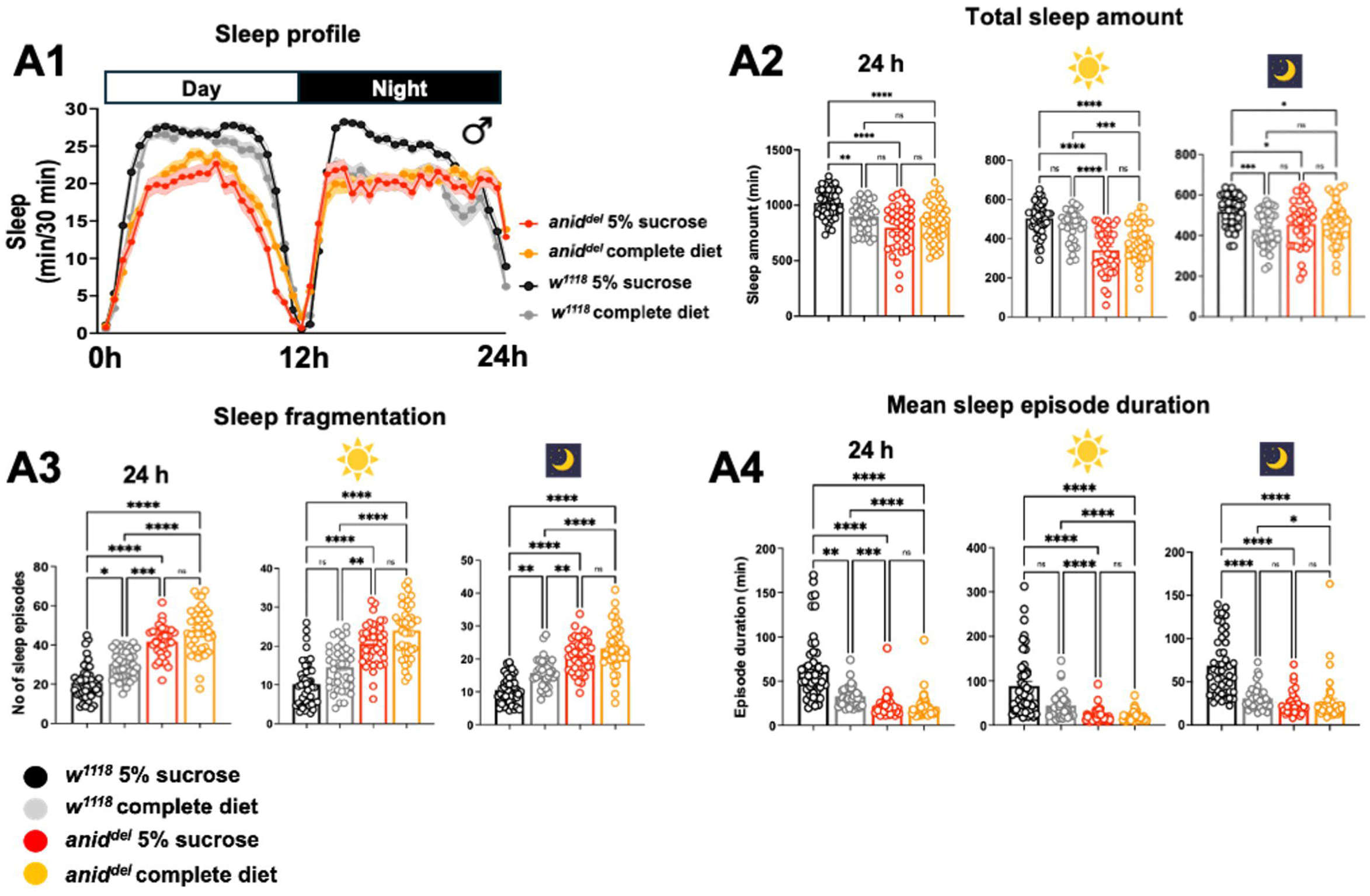
Diet-dependent modulation of sleep is impaired in *anid^d^*^e*l*^ mutant flies. **(A1-A4)** Sleep is reduced and fragmented in *w^1118^* on complete diet compared with 5% sucrose diet, whereas *anid^del^* mutants fail to show this diet-dependent modulation. (A1) Sleep profiles of *w^1118^* males on complete food (black; n = 39) and 5% sucrose food (gray; n = 47), and *anid^del^* males on complete food (orange; n = 41) and 5% sucrose food (red; n = 38). (A2) Total sleep (min). (A3) Number of sleep episodes. (A4) Mean sleep episode duration (min). All parameters were analyzed over 24 h and during daytime and nighttime periods separately. Data are presented as mean ± SEM. Statistics as described in Figure 2. See also Figure S5.

### ANIDRA regulates mTOR activity in glia

The mTORC1 (mechanistic Target of Rapamycin Complex 1) pathway is central to nutrient sensing and adjust metabolism, physiology and behavior accordingly^48^. Thus, we wondered whether ANID impacts mTOR activity in glial cells. First, we used the transcriptional M-box-GFP reporter (nuclear), which is used to determine mTOR C1activity in glia^49^. GFP levels are inversely correlated with mTORC1 activity and are thus elevated when animals experience dietary restrictions. Indeed, we found that control flies showed higher 4M-box activity on sucrose-only food. Strikingly, in *anid^del^* mutants, 4M-box activity was constantly low, indicating constitutive mTORC1 activation (Figure 6A, 6B, S6A). We corroborated these observations by measuring levels of ATG8 a key autophagy effector, whose expression is suppressed by active mTORC1. Indeed, in mutant flies, ATG8 levels were suppressed on sucrose-only food, further indicating elevated mTORC1 activity (Figure 6SB, 6SC).

**Figure 6.**
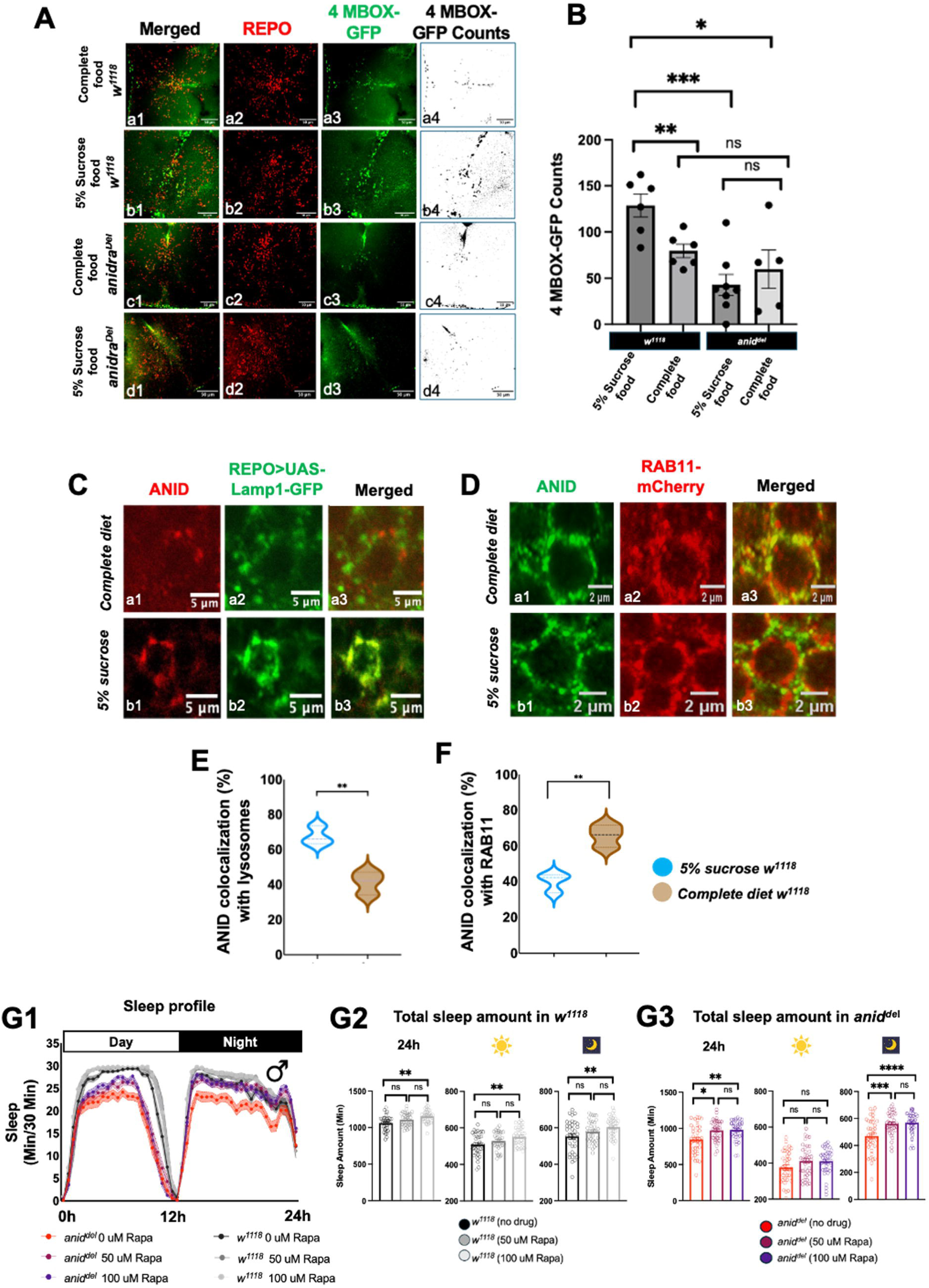
Diet-dependent mTORC1 signaling is altered in glia of *anid^del^* mutants. (A) mTORC1-dependent MITF (microphthalmia-associated transcription factor) transcriptional activity in glia of *w^1118^* and *anid^del^* male flies. Flies expressing the mTORC1-repressed reporter 4MBOX-GFP were maintained on complete or 5% sucrose food for 5 days, and brains were immunostained with anti-GFP and anti-REPO. Rows a-d represent (a) *w^1118^* on complete food, (b) *w^1118^* on 5% sucrose food, (c) *anid^del^* on complete food, and (d) *anid^del^* on 5% sucrose food. Columns 1- 4 represent: merged image, anti-REPO (red), anti-GFP (green), and colocalization of REPO and GFP channels used for quantification of nuclear puncta reflecting 4MBOX-GFP expression in glial nuclei (ImageJ), respectively. (B) Quantification of 4MBOX-GFP nuclear puncta in brains from flies maintained on complete food (*w^1118^*, n = 6; *anid^del^*, n = 5, males) or 5% sucrose food (*w^1118^*, n = 6; *anid^del^*, n = 8, males). (C) Colocalization of ANID with the lysosomal marker LAMP1-GFP expressed in glia under *repo*-*GAL*4 control. Rows a and b represent *w^1118^* flies on complete and 5% sucrose food, respectively. Columns show anti-ANID (red; a1, b1), anti-GFP detecting LAMP1-GFP (green; a2, b2). (D) Colocalization of ANID with the recycling endosome marker RAB11-mCherry expressed in glia under *repo*-*G*AL*4* control. Rows a and b represent *w^1118^* flies on complete food and 5% sucrose food, respectively. Columns show anti-ANID (green; a1, b1), anti-mCherry detecting RAB11-mCherry (red; a2, b2), and merged images (a3, b3). (E) Quantification of ANID colocalization with LAMP1-GFP in lysosomes of glia on 5% sucrose food (n = 5) versus complete food (n = 5). (F) Quantification of ANID colocalization with RAB11-mCherry in glia on 5% sucrose food (n = 9) versus complete food (n = 9). **(G1- G3)** Rapamycin (Rapa) treatment improves sleep in *anid^del^*mutants. Sleep behavior in *anid^del^* males maintained on 5% sucrose food alone (red; n = 38, no drug), 5% sucrose + 50 µM rapa (magenta; n = 39), or 5% sucrose + 100 µM rapa (purple; n = 39), and *w^1118^* males maintained on 5% sucrose food alone (black; n = 36, no drug), 5% sucrose + 50 µM rapa (gray; n = 37), or 5% sucrose + 100 µM rapa (light gray; n = 36). (G1) Sleep profiles. (G2- G3) Total sleep (min). Sleep duration analyzed over 24 h and during daytime and nighttime periods separately. Scale bars for images are indicated in each panel. Data are presented as mean ± SEM. Statistical comparisons for panels B, C1-C3 as described in Figure 2. Statistics for panels F, and G were performed as described in Figure 1.

Members of the PAT family of transporters can function as “Transceptors”, membrane proteins that function simultaneously as a nutrient transporter and as a signaling receptor. Upon binding with amino acids, they change their configuration and modulate the activity of targets such as mTOR^50^. We therefore wondered whether ANID and mTOR colocalize in subcellular compartments. Indeed, we found that this was the case: ANID and myc-tagged mTOR showed broad colocalization inside Cortex and Ensheathing glial cells (Figure S6E). Moreover, ANID subcellular localization changed as a function of diet. ANID colocalization with the recycling endosomal marker RAB11 was increased under complete diet (Figure 6D, 6F), while under sucrose-only diet ANID was preferentially localized in the lysosome, a major site for mTOR activity (Figure 6C, 6E). Combined with the constant activation of mTORC1 in *anid^del^* mutants, these results strongly suggest that under sucrose-only diet, ANID relocalizes from RAB11-positive endosomes to lysosomes to inactivate mTORC1.

To determine whether excessive mTORC1 activation contributes to the *anid^del^* sleep phenotypes, we exposed these mutants and wild-type flies to 50 or 100 μM rapamycin, a specific mTORC1 inhibitor. Sleep amount was weakly increased in control flies both during the day and night (Figure G1-3), which thus does not have a strong sedative effect on flies. A similar, though not significant, increase was observed with *anid^del^* flies during the day. However, at night, the *anid^del^* sleep profile and sleep duration were markedly improved (Figure G1-3, SE1-2), resembling those of wild-type flies. This nighttime specific effect of rapamycin is particularly interesting, since nighttime sleep is under much stronger diet-dependent modulation than daytime sleep (Figure 5).

Nighttime sleep fragmentation was mildly reduced in mutant flies in the presence of rapamycin (Fig S6E3-6). Exposure to rapamycin thus attenuates, but does not fully correct the *anid^del^* sleep phenotypes. This incomplete rescue might reflect the systemic nature of oral rapamycin administration, which affects mTORC1 broadly across tissues and cell types, rather than selectively within glia. Combined, these results support the model that ANID modulates sleep at least in part via mTORC1 signaling (see also discussion).

### ANID modulates the activity of amino acid sensitive wake promoting neurons regulating sleep

Neurons expressing the DH44 peptide in the Pars Intercerebralis (the insect homolog of the hypothalamus) are known to respond to dietary amino acids^30^. They are also critical circadian output neurons and promote arousal^31,32,51,52^. We therefore asked whether their activity was impacted by ANID. To probe DH44+ neurons activity in vivo, we expressed the ratiometric calcium sensor CAMPARI^53^ in these cells and measured their neuronal activity at two different time points (Zeitgeber [ZT] 1 and 7, i.e. 1 and 7 hours after the light-on transition) and two dietary conditions (complete vs sucrose-only diet). In wild-type animals, neuronal activity was low in the morning and elevated in the early afternoon^54^, and was further increased upon exposure to an amino acid containing diet, as previously described^30^. Indeed, we observed increased activity in wild-type flies at ZT7 compared to ZT1. Remarkably, in *anid^del^* mutants, DH44+ neurons showed persistently elevated activity, regardless of time-of-day or dietary condition (Figure 7 A-C). This indicates that ANID represses arousal neurons sensitive to amino acid under nutrient-poor conditions to control the sleep/wake cycle. Since DH44 neurons function downstream of the circadian neural network, their constant activation probably explains the increased arrhythmicity observed under DD (Figure S1A3).

**Figure 7.**
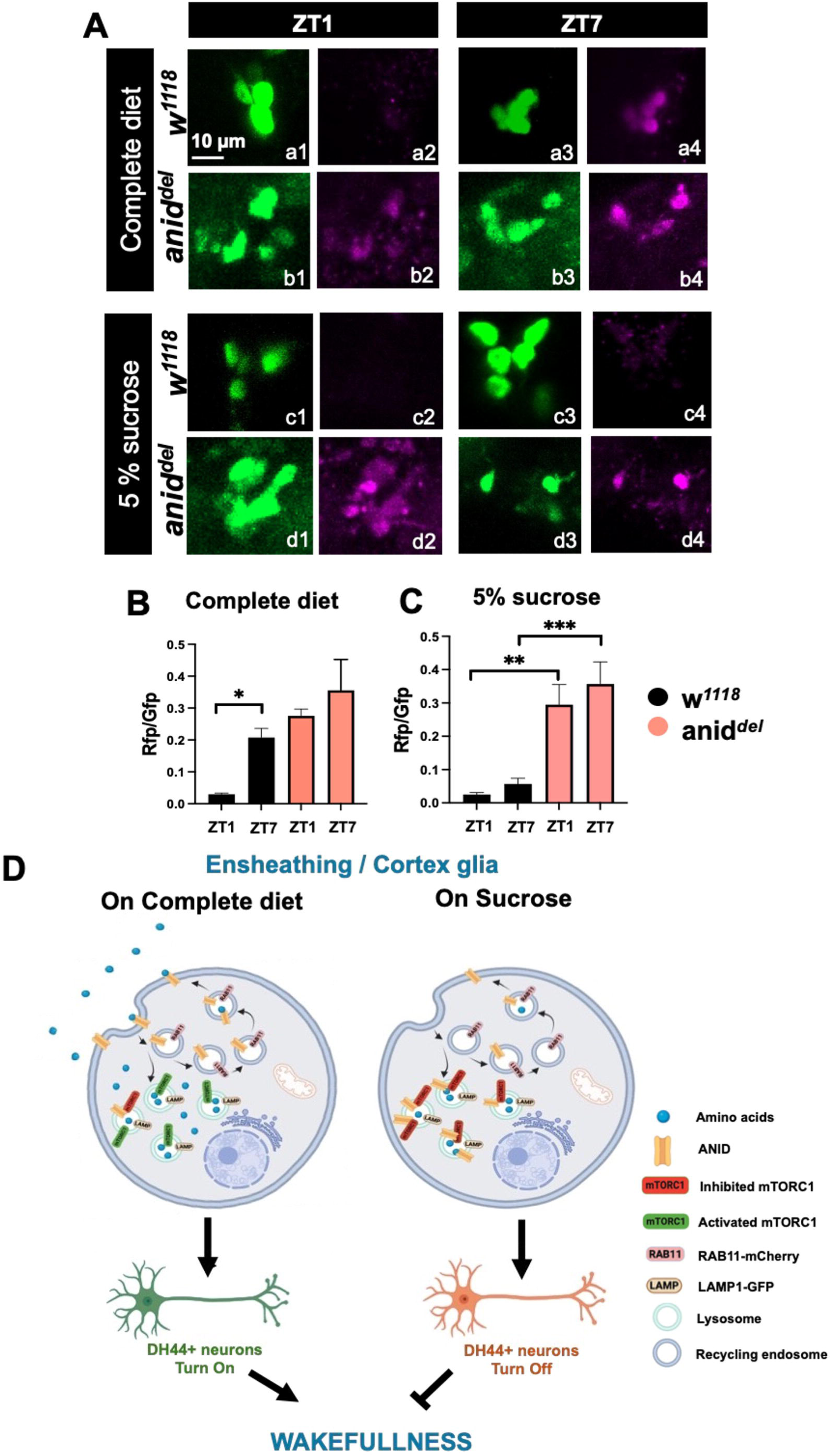
Loss of ANIDRA constantly activates amino acid-sensitive DH44 neurons. **(A-C)** CaMPARI-based imaging of DH44+ neuronal activity in *w^1118^* and *anid^del^* adult fly brains. **(A)** CaMPARI fluorescence in *DH44-GAL4 > UAS-CaMPARI* brains from *w^1118^* and *anid^del^* flies following 3-4 days of entrainment on complete or 5% sucrose food, imaged at ZT1 and ZT7 circadian time. Rows a-d represent (a) *w^1118^* on complete food, (b) *anid^del^* on complete food, (c) *w^1118^* on 5% sucrose food, and (d) *anid^del^* on 5% sucrose food. Within each row, columns 1- 2 show green (unconverted CaMPARI) and magenta (photoconverted CaMPARI) channels at ZT1, and columns 3- 4 show the same channels at ZT7. **(B)** Quantification of the CaMPARI red/green fluorescence ratio in DH44 neurons of *w^1118^* (n = 6) and *anid^del^* (n = 5) flies maintained on complete food at ZT1 and ZT7. **(C)** Quantification of the CaMPARI red/green fluorescence ratio in DH44 neurons of *w^1118^* (n = 5) and *anid^del^* (n = 5) flies maintained on 5% sucrose food at ZT1 and ZT7. **(D)** Proposed model for glial ANID function in the regulation of sleep. Scale bar for images is indicated in the panel. Data are presented as mean ± SEM. Statistics as described in Figure 1.

## Discussion

Diet and sleep are intimately related^16–18,55^. Indeed, animals can either rest or search for food. The nutrient composition of the diet plays a critical role in this choice, as animals need to consume a balanced diet that allows to replenish their stores of sugar, fat and amino acids to ensure proper bodily functions. How diet determines sleep has been the focus of a great number of studies in humans and animal models^16,18,55–57^, but a detailed understanding of the mechanisms by which diet determines sleep amount and quality is still lacking. Our work addresses this gap, revealing that the amino acid transporter ANID plays a central role in allowing *Drosophila* to adjust sleep amount and quality as a function of diet. Indeed, we found that ANID promotes sleep when flies are only fed sucrose. In its absence, flies behave as if they are constantly on a nutritionally balanced diet rich in amino acids: total sleep duration is decreased and more fragmented in the *anid* mutants, as observed in wild-type flies when they are on a complete diet. We thus surmise that ANID functions as an emergency brake in case of severe amino acid depletion, inhibiting the mTORC1 pathway and the activity of DH44 neurons, which promote arousal, as well as feeding. This results in increased rest on poor diet, which might help animal preservation of energy under nutritional hardship. Surprisingly however, ANID mutants do not detectably consume more food in the days during which sleep is being measured, even though DH44+ neurons activation was previously shown to be sufficient to increase feeding^30^. It is possible that ANID modulates additional neurons that repress food consumption, at least temporarily. Alternatively, long-term activation s of DH44+ neurons may reduce their impact on feeding, for example through satiety feedback.

Our work and a previous study^17^ found that a diet rich in amino acids reduces sleep. Yet, when taken in isolation, certain amino acids actually promote sleep in *Drosophila* (L-threonine, L-arginine, L- histidine, D-serine and D-glutamine), while others have the opposite effect (L-phenylalanine, L-leucine)^26,27^. Since *anid* mutants behave as if they are constantly on nutrient-rich diet, we cannot simply supplement amino acids to sucrose-only food to identify amino acids that ANID senses *in vivo* to regulate sleep. However, our results in HEK293 cells give us some interesting clues. Of the four amino acids we tested for transport, ANID could transport them all. Those belongs to very different categories of amino acids, suggesting that ANID has a broad substrate specificity which has been observed for other members of this protein family^58^. Transport efficiency was strongly impacted by pH, with pH6.5 appearing to be optimal. Interestingly, this is the pH of RAB11-positive recycling endosomes, where ANID resides preferentially under complete food. Sodium also impacted transport for specific amino acids, thus providing a potential additional level of activity modulation based on ANID’s subcellular location. Competition experiments further refined our understanding of the type of amino acids ANID can interact with. Most amino acids we tested (9 out of 16) significantly reduced proline transport, further supporting the notion that ANID is a broad-specificity transporter. Among them were the two acidic amino acids (L-glutamate and L-aspartate), along with L-alanine. This is interesting, because the transporter CG13248 permits DH44+ neurons to sense L-glutamate, L-aspartate and L-alanine^30^. These three amino acids might thus be particularly important to modulate diet-dependent behaviors, such as feeding and sleep. They might acutely promote food consumption by directly activating DH44 neurons through CG13248, while having long-term effects on sleep via ANID, which in turn regulates DH44 activity and thus shaping sleep/arousal dynamics differently.

Our study also reveals the key role ensheathing and cortex glia play in regulating diet-dependent sleep. We note that another amino-acid transporter expressed in ensheathing glia, EAAT2, has been proposed to regulate sleep by interacting with taurine, a sleep-promoting amino acid that is not incorporated into protein synthesis^42^. However, taurine is a GABA receptor agonist, and thus likely promotes sleep through this receptor, rather than through the mTOR pathway. EAAT2 would regulate taurine’s availability through its transport activity. Interestingly, the amino acid transporters JHL21 and MND are also implicated in sleep control, but their role is distinct from the ANID in multiple ways^28^. They function specifically in the blood-brain barriers (perineurial and sub-perineurial glia) and seem to play an important role in determining the availability of certain amino acids in the brain, such as leucine. Also, their transport activity promotes sleep rather than inhibiting it. Sleep modulation by amino acids is thus quite complex, with opposing effects of different glial subsets and transporters on sleep. Through differential amino acid specificity, this intricate regulation may enable files to fine- tune their sleep according to their precise metabolic status. In the same study^28^, the mTOR pathway was proposed to play a role in sleep in the blood brain barrier. However, the evidence did not directly associate JHL21/MND and their regulator BLOC1 to mTORC1 signaling. Our work directly connects this pathway to ANID. Indeed, in the absence of ANID, mTORC1 is constantly activated, and suppressing this pathway partially rescue the ANID mutant sleep phenotype. There are several reasons that could explain a partial rather than a full rescue after exposure to the mTORC1 inhibitor rapamycin.

First, rapamycin may not reach with sufficient efficiency to ensheathing and cortex glia to completely suppress mTORC1. Second, the complex and multi-tissue function of mTORC1 in sleep regulation could prevent us from observing a complete suppression of the ANID phenotype through pharmacological intervention. Finally, ANID loss may engage additional signaling pathways within glial cells that operate independently of mTORC1. However, it is striking that in mutant animals, rapamycin improved specifically night sleep, which is most sensitive to diet in our experimental conditions.

In summary, our work reveals a novel and critical pathway by which fruit flies sense the quality of their diet and thus adjust sleep accordingly. We propose that ANID functions as a transceptor, like other members of the PAT1 family of transporter^50^. Upon binding to amino acids, ANID would undergo a conformational change and translocate from the lysosome to RAB11+ vesicles. This translocation would result in the increase of mTORC1 signaling, and the activation of arousal promoting neurons such as those expressing DH44 (Figure 7D). Conversely, ANID could translocate from RAB11+ vesicles to the lysosomes when amino acids concentration drops. How glial cells stimulate DH44+ or other neurons implicated in sleep/arousal regulation will need to be studied in the future. It will also be interesting to determine whether this glia to sleep/arousal neuron connection is a conserved mechanism of sleep modulation.

## Acknowledgments

We are very grateful to A. Nicole Fox for sharing unpublished data identifying *anid* as an astrocyte-enriched mRNA. We thank Michael Brodsky and the University of Massachusetts Medical School Mutagenesis Core for assistance with the design and generation of the Anidra CRISPR gRNA vector. We are also thankful to Niraj Nirala for his help with gut dissection and staining. We thank Lauren North for help with the Café assay, and Vinh Phan for technical support. Finally, we thank the labs of N. Perrimon, G. Emery and E. Baehrecke, as well as the VDRC and Bloomington stock centers for fly strains. This work was funded by the National Institute of General Medical Sciences (1R35GM145253 to PE), the National Institute on Drug Abuse (R01DA035224 to HEM) and the National Institute of Neurological Disorders and Stroke (NS124146 and NS053538 to MRF)

## Author Contributions

Conceptualization, R.C. and P.E.; methodology, R.C., R.R.F., C.C, T.S., M.R.F., H.E.M and P.E..; Investigation, R.C., R.R.F., C.C and T.S; writing – original draft, R.C. and P.E.; writing – review & editing, R.C., R.R.F.,C.C, T.S., M.R.F., H.E.M and P.E, funding acquisition, P.E. H.E.M. and M.R.F.; supervision, P.E. H.E.M. and M.R.F.

## Material and Methods

### Fly strains and rearing conditions

Prior to behavioral monitoring or other experiments, all flies were reared on standard cornmeal/agar medium supplemented with yeast. The following *Drosophila* strains were used: *w^1118^* (Vienna *Drosophila* Stock Center); *4MBOX-GFP*^49^ (gift from Norbert Perrimon lab, Harvard Medical)*; UAS-Rab11-mCherry*^59^ (gift from Gregory Emery, University of Montreal)*; UAS-ATG8-GFP* (BDSC 52005 gift from Eric Baehrecke, UMass Chan Medical School)*; repo- GAL4, alrm-GAL4. mz0709-GAL4, np2222-GalGAL4. UAS-mCD8::GFP* (BDSC 5137), *UAS-Cas9* (BDSC 58985), *DH44-GAL4* (BDSC 51987, check on vial)*, UAS-CaMPARI* (BDSC 78317), *UAS-Lamp1-GFP* (BDSC 42714, this strain also carries nSyb-*GAL4* on 3^rd^ chromosome, which was removed for our experiments), and *UAS- TOR*-*myc* (BDSC 53727) were obtained from the Bloomington Stock Center, USA.

The following strains were generated for this study: *anidra* (*CG7888*) mutant (*anid^del^*), *UAS-anid, and UAS-anid^gRNA^. anid^del^*flies were backcrossed into the *w^1118^* genetic background for 6 generations. Note that *anid^del^* flies require regular backcrossing, as they tend to accumulate genetic modifiers that suppress their total sleep duration phenotype. The sleep fragmentation phenotypes appear largely insensitive to this genetic drift.

### CRISPR/Cas9-guided generation of the *anid^del^* mutant line

The *anidra* specific guides GGATTGTAGTCGGGATCCT and CGGCACGTATACCTCACTCA were inserted into the vector pCFD4^60^. The resulting construct was inserted into the *Drosophila* genome via phiC31-mediated site directed integration at the landing site attP40 using standard procedures (Rainbow Transgenics). To generate CRISPR deletions *actin5C*-Cas9 females were crossed to the *anidra* guide insertion. The resulting male founder animals were outcrossed to a TM6B balancer stock. 15 pools of 8 founder males carrying actin5C-Cas9 and the CG7888 guide insertion were screened for big deletions in the *anidra* locus by PCR using the primers *CG7888-forward* (AATCAGAAAGGGGAAAACGTCG) and *CG7888-reverse* (CGGATCTCAGCTCAACTGTAAC). Subsequently, individual male progeny from several positive founder pools were backcrossed to a TM6B stock. After 3- 4 days, individual males were removed and again tested for newly induced large deletions between the two CRISPR sites by genomic PCR. For positive events stable stocks were established. Finally, a homozygous viable deletion mutant was then outcrossed to a *w^1118^* wild type stock for six generations to remove potential background hits.

### Generation of *UAS-anid*

*UAS-anid* was generated by amplifying the ORF from cDNA GH9436 (Berkeley Drosophila Genome Project) using the primers CG7888-NotI-for (ACTGATAGCGGCCGCcaaaacATGACCAAGAATGGACACAGCAAT) and CG7888-XbaI-rev (ccCATCGCtctagaCTTGAATTGGAATGCCCGCT) and inserting the amplicon into the pUAST vector using NotI and XbaI restriction sites. Transgenes were generated at BestGene Inc (Chino Hills, CA).

### Generation of *UAS-anid^gRNA^*

Guide RNAs (gRNAs) targeting *cg7888* in *Drosophila melanogaster* were designed using CRISPR target prediction tools (flyCRISPR) to minimize off-target effects. Two gRNAs were selected to generate a deletion allele by targeting two distinct regions within the genomic locus. The target sequences were:

gRNA-1: GGCTTACGTCGCCGATCATC gRNA-2: ATATGCAAACTACTCCGAAA

Complementary sense and antisense oligonucleotides containing cloning overhangs were synthesized, annealed, and ligated into a pCFD6 vector (Addgene #73915) following standard cloning procedures. The resulting constructs were verified by Sanger sequencing and inserted into the second chromosome using phiC31 integrase by the BestGene Inc (Chino Hills, CA).

### ANIDRA antibody generation

A custom-made polyclonal antibody against ANIDRA was generated by 21st Century Biochemicals. A synthetic peptide corresponding to the ANIDRA amino acid sequence **MTKNGHSNSAYVADHPDKLC-amide** was used as the immunogen. Peptide synthesis was performed in an ISO 9001:2015-certified peptide manufacturing facility, followed by HPLC purification to approximately 85% purity. Peptide identity and sequence were confirmed by mass spectrometry, including tandem MS (CID MS/MS), nanospray MS, and HPLC analyses. The peptide was conjugated to an immune carrier protein using m-maleimidobenzoyl-N-hydroxysuccinimide ester (MBS) chemistry prior to immunization. Polyclonal antibody production was carried out in two rabbits according to the company’s standard immunization protocol.

### Sleep measurement and analysis

Sleep experiments were performed using adult flies aged 3- 5 days. Individual flies were placed in 5 × 65 mm glass tubes containing 2% agar and 5% sucrose and loaded into *Drosophila* Activity Monitors (DAM; Trikinetics, Waltham, MA, USA) (https://www.trikinetics.com/). Locomotor activity was recorded at 1- minute intervals.

Experiments were conducted in I36LL incubators from Percival Scientific under a 12 h:12 h light- dark (LD) cycle at 25°C. Flies were maintained under these conditions for 6- 7 days, and sleep analysis was performed using data averaged from days 4- 6.

Data were collected using DAMSystem3 and DAMFileScan111 software ((https://www.trikinetics.com/), and processed with the Sleep and Circadian Analysis MATLAB Program (SCAMP) developed in MATLAB^61^. A period of at least 5 minutes of inactivity was defined as sleep. Sleep parameters analyzed included sleep profile (sleep time per 30 min), sleep frequency (number of sleep bouts during day and night), mean sleep bout duration (min), pDOZE (probability of falling asleep), pWAKE (probability of waking up), and mean activity per min while awake.

### Circadian behavior

Male flies were entrained to a 12 h:12 h light–dark (LD) cycle at 25°C for 3 days and then transferred to constant darkness (DD) for 6 days. Locomotor activity was monitored using the DAM system, as described in the sleep experiment section.

Circadian period length and rhythmicity were analyzed using the FAAS-X software developed by M. Boudinot and F. Rouyer (Centre National de la Recherche Scientifique, Gif-sur-Yvette Cedex, France)^62^. Actograms and eduction graphs were also generated using FAAS-X.

### Mechanical sleep deprivation assay

Sleep deprivation experiments were carried out in a Percival Scientific incubator under a 12 h:12 h light- dark (LD) cycle at 25°C. Mechanical sleep deprivation was induced using a customized vortexer system (VMP Vortexer Mounting Plate, Trikinetics, Waltham, MA) placed on a cushion to minimize vibration within the incubator. Briefly, individual flies were housed in glass tubes and loaded into Drosophila Activity Monitors (DAMs), which were connected to the vortexer system for sleep deprivation.

Flies were first habituated for 3 days, followed by 2 days of baseline sleep recording. Mechanical sleep deprivation was then applied during the 12- hour night, after which flies were allowed to recover for 48 hours under the same LD conditions. During deprivation, the monitors were shaken for 2 seconds at random intervals within every 20- seconds window throughout the night using a software-controlled program from Trikinetics. Sleep and activity data were analyzed using SCAMP.

### CAFÉ assay

To measure food consumption in *anid^del^* mutants and *w¹¹¹* flies, the CAFÉ assay was performed as described by Ja et al.^47^with some modifications to minimize evaporation. Briefly, empty *Drosophila* food vials were filled with 8 mL of distilled water at the base to maintain humidity. Foam plugs were cut into half with a razor blade, inserted into each vial, and positioned 3 cm from the top of the tube. A cap was tightly fitted onto each vial, with the 4 holes where a 200 µL plastic pipette tips can be inserted. Glass capillaries (Wiretrol® II, cat# 5-000-20-20, from Drummond scientific company) were threaded through trimmed 200 µL plastic pipette tips, secured with a snug friction fit, and loaded with 5% sucrose solution containing to the 20 µL calibration mark. Four such capillary assemblies were inserted through each vial cap, with all capillaries verified to be filled to the same 20 µL starting volume. Eight adult flies were introduced per vial, and the assembled setup was placed in an incubator under controlled conditions (25°C, 12h:12h LD cycle, 65% relative humidity). Inverted empty food vial was placed to the top of the setup to further minimize evaporation. Vials without flies served as evaporation controls. After 48 hours, meniscus displacement from the 20 µL baseline was recorded, and food consumption per fly was calculated by subtracting the control evaporation from the experimental values.

### Lifespan assay

For lifespan assay on different food (complete vs sucrose diet), 3- 4 days old adult flies were collected and maintained under standard laboratory conditions at 25°C under a 12 h light:12 h dark cycle. Flies were transferred to fresh food vials every 2 days and the number of dead flies was recorded at each transfer until all *anid^del^*flies had died.

Survival data were analyzed using Kaplan- Meier survival curves, and statistical significance between groups was determined using the log-rank (Mantel- Cox) test.

### Brain dissection and immunostaining

Fly heads were removed from the body and brains were dissected in PBST buffer (1× PBS containing 0.1% Triton X-100). Dissected brains were fixed in 4% paraformaldehyde for 20 min and then washed five times with PBST, followed by three washes in PBS for 5-10 min each. Samples were blocked in the blocking solution (10% normal donkey serum in PBST) for 2 h at room temperature and then incubated with primary antibodies for overnights at 4°C. After primary antibody incubation, brains were washed six times for 10 min each in PBST and then incubated overnight at 4°C with the appropriate secondary antibodies. Samples were again washed six times in PBST, three times in PBS and mounted in Vector Laboratories Vectashield antifade mounting medium.

Primary antibodies used included chicken anti-GFP (ab13970, 1:1000; Abcam), rabbit anti-ANID (1:500; generated in this study), mouse anti-Repo (DSHB 8D12, 1:800), mouse anti-GFP (MA5-47387, 1:1000; Invitrogen), and mouse anti-mCherry (NBP1-96752SS, 1:1000; Novus Biologicals). Secondary antibodies included Alexa Fluor 488 or Alexa Fluor 647-conjugated antibodies raised against chicken, mouse, or rabbit IgG (Invitrogen), all used at 1:1000 dilution. Images were acquired using a Zeiss LSM700 or Zeiss LSM800 confocal microscopes. Z-stack images were collected to examine protein localization, and image processing and analysis were performed using ImageJ (Fiji software).

### 4MBOX-GFP puncta counts

To quantify 4MBOX-GFP puncta co-localized with Repo, images were analyzed using ImageJ/Fiji. Co-labeled puncta were first identified from merged channel images within defined regions of interest (ROIs). Images were then converted to grayscale, and a consistent threshold was applied across all samples in the same experiment to separate puncta from background signal. Quantification of puncta was performed using the *Analyze Particles* function with appropriate size and circularity settings to minimize background noise and exclude non-specific signals.

### Protein extraction and western blotting

20 fly heads were collected for each sample, homogenized, and lysed in RIPA buffer containing protease inhibitors cocktail (PMSF, leupeptin, pepstatin A, aprotinin). The samples were shaken at 4°C for 30 min and then spun at 13000g for 10 min to remove insoluble material. Without boiling, samples were mixed with SDS-PAGE sample buffer (100mM Tris, pH 6.8, 4.4% SDS, glycerol, 100mM DTT, and 0.04% bromophenol-blue) and resolved by SDS-PAGE. Western Blots were stained with rabbit anti-ANID and anti-Tubulin (both 1:10000). Secondary antibodies conjugated with HRP from Jackson Immuno Research were used at 1:10,000 dilutions.

### CaMPARI imaging and analysis

To monitor calcium activity in DH44⁺ neurons, *UAS-CaMPARI*; *anid^del^* flies were crossed with *DH44-GAL4, anid^del^* flies, generating progeny expressing CaMPARI in DH44⁺ neurons in the *anid* mutant background. Male flies were entrained under 12 h:12 h light-dark cycles at 25°C for 3- 4 days. Flies were collected at two time points during the fifth day of LD, in the morning (ZT1) and midday (ZT7).

Photoconversion was performed essentially as previously described^63^. In brief, flies were gently mounted on Petri dishes and exposed to ultraviolet light (395–405 nm, ∼10 mW/cm² for 5 min to induce calcium-dependent conversion of CaMPARI from green to red fluorescence. Brains were immediately dissected in ice-cold extracellular fly saline under dim red light to avoid unwanted photoconversion. Samples were imaged using a ZEISS LSM 800 confocal microscope. Images were analyzed using ImageJ. After background subtraction, green (F_green) and red (F_red) fluorescence intensities were measured in DH44⁺ neurons, and CaMPARI activity was quantified as the ratio of red to green fluorescence (F_red/F_green).

### Dietary composition of complete diet and sucrose diet

Sucrose diet: sucrose (50g/L) with agar (20g/L)

Complete diet: Agar (6.5 g/L), Yeast (63 g/L), Cornmeal (60g/L), Molasses (60ml/L), Acid mix (4ml/L), Tegosept (1.3h/L)

### Rapamycin feeding experiments

Rapamycin (from MedChemExpress [MCE] Cat # HY10219) was dissolved in ethanol at a concentration of 10 mM. It was then diluted in 5% sucrose + 2% agar food solution to concentrations of 50 µM and 100 µM. Behavior tubes were prepared with these solutions, or with sucrose food with only ethanol as a control.

### Gut dissection and staining

Adult flies aged 5-7 days were used for gut dissection^64^. Whole guts (15-20 guts per genotype) were dissected and fixed in 1× PBS containing 4% formaldehyde (Fisher Biosciences) for 2 h at room temperature, followed by washes in 1× PBS and permeabilization in 1× PBS containing 0.1% Triton X-100. Blocking and incubation with primary and secondary antibodies were performed in 1× PBS containing 0.5% BSA, 5% normal horse serum (Vector Laboratories), and 0.1% Triton X-100. The primary antibodies used were rabbit anti-ANID (1:500) and mouse anti-GFP (1:1000).

Secondary antibodies included goat anti-rabbit IgG conjugated to Alexa Fluor 555, and goat anti-mouse IgG conjugated to Alexa Fluor 488 (Invitrogen), used at dilutions of 1:2000 and 1:1000, respectively. Tissues were mounted in DAPI-containing mounting medium (1.5 μg/ml; VECTASHIELD, Vector Laboratories). Images were acquired using a Nikon spinning-disk confocal microscope and processed with ImageJ.

### HA-anidra cloning and expression in HEK293T cells

The full-length cDNA of *Drosophila* CG7888 (*anidra*) was amplified from the GH09436 cDNA clone obtained from the Drosophila Genomics Resource Center (DGRC) and subcloned into the mammalian expression vector pcDNA3.1(+) for expression in HEK293T cells. Both N-terminal HA-tagged and C-terminal HA-tagged CG7888 constructs were generated. Correct insertion and reading frame of all constructs were confirmed by DNA sequencing.

HEK-293T cells cultured in DMEM (as mentioned in next section) were initially transiently transfected with HA-tagged *anidra* constructs to assess transporter activity. Comparative analysis of the N-terminal and C-terminal HA-tagged constructs demonstrated that the N-terminal HA-tagged CG7888 construct exhibited stronger and more reproducible uptake activity. Therefore, all subsequent time-course and amino acid uptake assays were performed using the N-terminal HA-tagged construct.

For further functional characterization, stable HEK293T cell lines expressing the N-terminal HA-tagged CG7888 construct were generated, and uptake assays were performed as described below.

### [^3^H] substrate uptake assay

HEK293T cells stably expressing HA-ANIDRA were maintained in DMEM supplemented with 10% fetal bovine serum, 2mM L-glutamine, and 10^2^units/mL penn-strep at 37°C in 5%CO_2._ One day prior to assay, 7.5×10^4^ cells/well were seeded onto poly-D-lysine-coated 96-well plates. To assess substrate uptake, cells were washed twice with Krebs-Ringers solution (120mM NaCl, 4.7mM KCl, 2.2mM CaCl_2_, 1.2mM MgSO_4_, 1.2mM KH_2_PO_4_) buffered with either 10mM HEPES (KRH; pH 7.4) or 10mM MES (KRM pH 6.0 or 6.5, as indicated), and pre-incubated in the same buffer supplemented with 0.18% glucose, 30 min, 37°C. For sodium dependence studies, NaCl was substituted with choline chloride. Uptake was initiated by adding a 1/10^th^ volume of a 10X concentrated substrate solution, and proceeded for 10min, 37°C. Uptake was terminated by three rapid washes with ice-cold KRH or KRM buffer. Cells were solubilized in scintillation fluid and accumulated [^3^H] was determined by liquid scintillation counting in a Wallac MicroBeta scintillation plate counter. Non-specific uptake was defined in parallel, either in the presence of 10mM 5-HTP (competition studies), or in vector-transfected cells (kinetic studies), and was subtracted from total uptake to determine ANIDRA-specific uptake.

***Kinetics studies*:** The indicated substrates were added at the indicated concentrations. Specific [^3^H] uptake was converted to pmol substrate uptake based on a 45% counting efficiency. Total cellular protein/well was measured using the BCA protein assay, in parallel wells solubilized in 1% Triton-X-100, and velocities were calculated as pmol/min/mg protein. Kinetic curves (velocity vs. [substrate]) were generated in GraphPad Prism, and were fitted using the Michaelis-Menten equation:

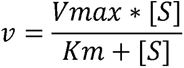

N values indicate biologically independent experiments; each performed with three technical replicates.

L-[2,3-^3^H]-alanine was obtained from American Radiolabeled Chemicals, and L-[2,3,4,5-^3^H]-proline and L-[3,4-^3^H]-glutamic acid were from Revitty.

### Surface biotinylation

Surface ANID was assessed by cell surface biotinylation as previously described by our laboratory^65–67^. Briefly, 1 x 10^6^ cells/well were seeded in poly-D-Lysine-coated 6-well plates one day prior to labeling, and surface proteins were covalently labeled by incubating twice x 15 min, 4°C in 1.0mg/mL sulfo-NHS-SS-biotin (Thermo Fisher) in phosphate-buffered saline supplemented with 0.1mM CaCl_2_ and 1 mM MgCl_2_ (PBS^2+^). Reactive NHS groups were quenched by washing thrice, and incubating 3 x 15 min, 4°C, in PBS^2+^ supplemented with 100mM glycine, and cells were lysed in RIPA buffer (10mM Tris, pH 7.4, 150mM NaCl, 1.0mM EDTA, 0.1% SDS, 1% Triton-X-100, 1% sodium deoxycholate) containing protease inhibitors (1.0mM PMSF and 1.0g/mL each leupeptin, aprotinin, and pepstatin). Lysates were cleared by centrifugation and protein concentrations were determined with the BCA protein assay (Thermo Fisher) using BSA prepared in RIPA buffer as a standard. Biotinylated proteins from equivalent amounts of lysate were isolated from non-biotinylated proteins by batch affinity chromatography using streptavidin agarose (Thermo Fisher) and were eluted with 2X Laemmli sample buffer by rotating, 30 min, RT. Eluted samples and a parallel amount of total lysate were resolved by 10% SDS-PAGE and proteins were transferred to nitrocellulose (Hybond ECL, Cytiva). HA-positive bands were identified by immunoblotting with high affinity rat anti-HA antibody (clone 3F10, Roche, 1:2000) and goat anti-rat secondary antibody conjugated to horseradish peroxidase (Jackson ImmunoResearch, 1:5000).

Immunoreactive bands were visualized by chemiluminescence using ECL Western Blotting substrate (Pierce/Thermo Fisher) and bands in the linear range of detection were captured using a ChemiDoc Imaging station (Biorad).

### Immunocytochemistry in HEK293T cells

HEK293T cells stably expressing HA-ANIDRA were seeded onto poly-D-lysine-coated glass coverslips in 24-well plates at a density of 2.5×10^4^ cells/well, 24 hours prior to assay. Cells were washed thrice in PBS and permeabilized in blocking solution (PBS, 5% normal goat serum, 1% IgG- and protease-free bovine serum albumin, 0.2% Triton-X-100), 30 min, room temperature. Cells were co-stained with high affinity rat anti-HA antibody (1:2000 in blocking solution) and either mouse anti-EEA1 (BDTransduction #610456), mouse anti-RAB11 (BDTransduction #610656) or mouse anti-RAB7 (D95F2, Cell Signaling Technology). 1 hour, room temperature, were washed thrice with PBS, and incubated 4 x 5 min in PBS, room temperature. Cells incubated with the indicated AlexaFluor-conjugated secondary antibodies (1:5000 in blocking solution, Jackson ImmunoResearch), 1 hour, RT, and were washed thrice in PBS. Coverslips dried at 37° and were mounted in ProLong Gold containing DAPI (Invitrogen). Cells were visualized with a Zeiss Axiovert 200M microscope using a 63×, 1.4 numerical aperture (NA) oil-immersion objective and 0.2 μm optical sections were captured through the *z*-axis with a Regita-R1 cooled CCD camera (Qimaging). 3D *z*-stack images were deconvolved with a constrained iterative algorithm using measured point spread functions for each fluorescent channel using Slidebook version 5.0 software (Intelligent Imaging Innovations). All representative images shown are single, 0.2μm planes through the center of each cell.

### Quantification and Statistical Analysis

GraphPad Prism 11 was used for graph generation and statistical analysis of independent datasets. Data are presented as mean ± S.E.M. unless otherwise stated in the figure legends. Normality was assessed using the D’Agostino- Pearson omnibus test. In general, for normally distributed (Gaussian) data, comparisons between two groups were performed using a two-tailed Student’s t-test, while comparisons among multiple groups were analyzed using one-way ANOVA followed by Tukey post hoc test, unless otherwise specified in the figure legends. For non-normally distributed data, the Mann- Whitney U test was used for comparisons between two groups, and the Kruskal–Wallis test followed by Dunn’s post hoc test was used for multiple-group comparisons. Statistical significance was defined as ****P < 0.0001, ***P < 0.001, **P < 0.01, and *P < 0.05. Specific statistical tests used for individual experiments are indicated in the corresponding figure legends.

---

**Figure S1. Sleep and circadian behavior in *anid^del^* mutant flies, related to Figure 1**

**(A1-A4)** Circadian locomotor behavior under 12h/12h LD conditions in *w^1118^* and *anid^del^* mutant male flies. White bars represent activity averages during the day, and dark bars during the night. (A1-2) Activity plots of *anid^del^*(n=14) and *w^1118^* (n=16), showing their average locomotor activity under a 12/12-hour (h) LD cycle, The morning (M) and evening (E) anticipatory behavior driven by the circadian clock are present in the mutants, (A3) Rhythmicity percentage (Fisher’s exact test), (A4) Circadian period lengths (two tailed t-test). N=32 fir *w^1118^* and N=29 for *anid^del^*

**(B1- B7)** Sleep behavior in *w^1118^* and *anid^del^* mutant female flies. (B1) Sleep profiles of *anid^del^* (magenta; n = 36) and *w^1118^* (wild type; black; n = 45) females. (B2) Total sleep (min). (B3) Number of sleep episodes. (B4) Mean sleep episode duration (min). (B5) pDOZE. (A6) pWAKE. (A7) Mean beam breaks per waking minute (locomotor activity).

All parameters were analyzed over 24 h and during daytime and nighttime periods separately.

**(C1- C2)** Quantification of sleep lost during sleep deprivation (SD). (C1) Total amount of sleep lost during SD (min). (C2) Percentage of total sleep lost during SD.

Data are presented as mean ± SEM. Statistics as described in Figure 1.

**Figure S2. ANIDRA expression in the optic lobe and additional sleep parameters following pan-glial *anid^del^*, related to Figure 2**

**(A)** ANID expression in the optic lobe of the adult *Drosophila* brain. CD8-GFP was expressed in astrocytic glia using *Alrm-GAL4*. Rows a, b, and c represent merged, anti-ANID (red), and anti-GFP (green) channels, respectively. (a1- c1) *anid^del^* mutant brain serving as a negative control. (a2- c2) Wild-type brain. White arrow indicates a region of strong ANID immunoreactivity that, based on morphology, is consistent with expression in cortex glia.

**(B)** Locomotor activity expressed as mean beam breaks per waking minute in control and glia-rescued *anid* mutants.

**(C)** *anid^de^*^l^ and *repo-GAL4>UAS-CD8-GFP* expression in hindgut (a1-a4) and midgut (b1-b4), (a1-b1) There is no GFP expression under *repo-GAL4* in enterocytes, (a2-b2), while ANID is expressed in hindgut enterocytes, (a3-b3) DAPI staining, (a4-b4) merged image. Pale autofluorescence is visible in the green and red channels.

Scale bars are indicated in each panel.

Data are presented as mean ± SEM. Statistics as described in Figure 2.

**Figure S3. Locomotor activity levels and severely reduced ANID levels following glial-specific CRISPR-mediated *anid* deletion, related to Figure 3**

**(A)** Locomotor activity after pan-glial *anid* deletion. Locomotor activity expressed as mean beam breaks per waking minute. Genotypes: *UAS-anid^gRNA^* (light magenta), *UAS-Cas9*; *repo*-*GAL*4 (magenta), and *repo*-*GAL4* > *UAS-Cas9*; *UAS-anid^gRNA^* (red) male flies.

**(B)** Locomotor activity after ensheathing glial deletion using *mz0709-GAL4*. Genotypes: *UAS-anid^gRNA^* (light magenta), *UAS-Cas9*; *mz0709*-*GAL4* (magenta), and *mz0709-GAL4* > *UAS-Cas9*; *UAS-anid^gRNA^* (red) male flies.

**(C)** Locomotor activity after cortex glial deletion using *np2222*-*GAL4*. Locomotor activity. Genotypes: *UAS-anid^gRNA^* (light magenta), *UAS-Cas9*; *np2222*-*GAL4* (magenta), and *np2222*-*GAL4* > *UAS-Cas9*; *UAS-anid^gRNA^* (red) male flies.

**(D)** Confocal images showing severely reduced ANID expression following pan-glial CRISPR-mediated *anid* deletion. Brains were stained with anti-ANID (green) and anti-REPO (red). (b1) *w^1118^* (b2) *anid^del^* mutant. (b3) Pan-glial CRISPR deletion using *repo*-*GAL4*.

**Figure S4. ANIDRA is a proton-activated amino acid transporter, related to Figure 4**

**(A)** *Cell surface biotinylation.* ANID plasma membrane localization was determined by cell-surface biotinylation in HEK293T cells stably expressing HA-ANID, as compared to vector-transfected control cells.

**(B)** ANID co-localizes with RAB11^+^ recycling endosomes in HA-ANIDRA-HEK293T cells. *Immunocytochemistry.* Cells were co-stained with anti-HA (red) and anti-endosomal compartment antibodies (green), and mounted in media containing DAPI (blue). (B1) EEA1, an early endosome marker; (B2) RAB11, a recycling endosome marker; (B3) RAB7, a late endosome marker. Scale bar = 20µm.

**(C)** 5HTP is a competitive ANIDRA inhibitor*. Proline uptake assay.* [^3^H]proline (100µM) uptake was measured in stably- transfected HA-ANIDRA HEK293T cells following pre- incubation with the indicated 5HTP concentrations, 30min, 37°C. Average data are presented as %specific proline uptake, defined with 20mM proline. Average 5HTP IC_50_ = 2.0±0.4 mM, n=2.

**(D-G)** ANIDRA can transport multiple amino acids and is differentially dependent on Na^+^. Saturation uptake kinetics for proline (D), glutamate (E), alanine (F) and glutamine

(G) were measured at pH 6.5, in the presence of Na^+^. K_m_ and V_max_ kinetic constants ±Na**^+^** are reported in Table I.

(H) Proline transport by ANIDRA is sensitive to multiple amino acids. *Amino Acid Competition Assay.* Cells co-incubated with 20mM of the indicated amino acid and 100µM [^3^H]proline in a Na^+^-containing buffer at pH 6.5. Average data are presented as %specific proline uptake ±S.E.M, defined with 10mM 5HTP. **p<0.01, *p<0.05, significantly less proline transport than (-)inhibitor control, one-way ANOVA (p=0.005) with Dunnett’s multiple comparison test.

**Figure S5. Survival and food consumption of *anid^del^* mutant flies on complete and sucrose diets, related to Figure 5**

**(A- C)** Survival curves of *w^1118^* and *anid^del^* and glia rescue of *anid* flies on complete food and 5% sucrose food.

(A) Survival of *w^1118^* (n = 60) and *anid^del^* (n = 65) male flies on 5% sucrose food.

(B) Survival of *anid^del^* male flies maintained on complete food (n = 55) or 5% sucrose food (n = 65, same flies as in A).

(C) Survival of *anid^del^* male flies with pan-glial rescue of *anid* expression.

Genotypes: *repo*-*GAL4* > *UAS-anid*; *anid^del^* (red; n = 45), *repo*-*GAL4*, *anid^del^* (purple; n = 50), *UAS-anid*; *anid^del^* (magenta; n = 50), *repo*-*GAL4*, *anid^del^*/+ (gray; n = 38), and *UAS-anid*; *anid^del^*/+ (black; n = 51).

**(D- E)** Food intake in *w^1118^* and *anid^del^* male flies measured by CAFE assay.

(D) Food consumption after 7 days on complete food in *w^1118^* (n = 24) and *anid^del^* (n = 24) male flies.

(E) Food consumption after 4 days on complete food followed by 2- 3 days on 5% sucrose food in *w^1118^* (n = 32) and *anid^del^* (n = 32) male flies.

(F) Representative image of the swollen abdomen phenotype observed in *anid^del^* mutant flies maintained on 5% sucrose food for 17- 20 days.

Data are presented as mean ± SEM. Statistical comparisons for the CAFE assay were performed as described in Figure 1. Survival curves were compared using the log-rank (Mantel-Cox) test. Different alphabets indicate significant difference.

**Figure S6. Food-dependent alterations in glial mTORC1 signaling and autophagy in *anid^del^* mutants, related to Figure 6**

**(A)** 4MBOX-GFP reporter activity reflecting mTORC1 signaling in brains of *w^1118^* and *anid^del^* adult male flies. Flies were maintained on complete food or 5% sucrose food for 5 days, and brains were immunostained with anti-GFP (white). Row-a shows whole-brain images; row-b shows higher-magnification views of the central brain, with the central complex (CC) outlined by a circle and the subesophageal zone (SEZ) indicated by a white arrow. Columns 1- 4 represent (1) *w^1118^* on complete food, (2) *w^1118^* on 5% sucrose food, (3) *anid^del^* on complete food, and (4) *anid^del^*on 5% sucrose food. Yellow box in the full brain indicates the region of interest including CC and SEZ.

**(B)** ATG8 puncta in ensheathing and cortex glia of *w^1118^* and *anid^del^* male adult flies maintained on 5% sucrose food. ATG8 (green) was expressed selectively in glia using *repo-GAL4* > *UAS-atg8* as an autophagy reporter. Row-a shows *w^1118^* and row-b shows *anid^del^*; columns 1 and 2 show ensheathing glia and cortex glia, respectively. (a1) Ensheathing glia, *w^1118^*. (a2) Cortex glia, *w^1118^*. (b1) Ensheathing glia, *anid^del^*. (b2) Cortex glia, *anid^del^*.

**(C)** Quantification of mean overall ATG8 fluorescence intensity in glia of *w^1118^* (black) and *anid^del^* (red) flies on 5% sucrose.

**(D)** ANID and mTOR-myc colocalization in the glia of adult *w^1118^* fly brain fed on 5% sucrose. (a1) ANID immunostaining (green), (a2) myc immunostaining (red) in brain expressing *UAS- TOR-myc* under *repo-GAL4*, (a3) merged.

**(E1- E6)** Rapamycin (Rapa) treatment in *anid^del^*mutants and *w^1118^* as described in Figure 6 (G1-G3). (E1 and E2). Same sleep profiles as in Figure 6G1, but mutant and wild-type traces are plotted separately to emphasize the robust improvement of nighttime sleep in the mutant animals fed rapamycin. (E3) Number of sleep episodes. (E4) Mean sleep episode duration (min), (E5) pDOZE, (E6) pWAKE, All parameters were quantified over 24 h and further subdivided into daytime and nighttime periods.

Scale bars for images are indicated in each panel. Data are presented as mean ± SEM. Statistics as described in Figure 1.

