## Supplementary figures and images for "Diet-dependent sleep modulation by the *Drosophila* amino acid transporter ANIDRA"

### Supplemental Figures 1-6

#### Slide 1
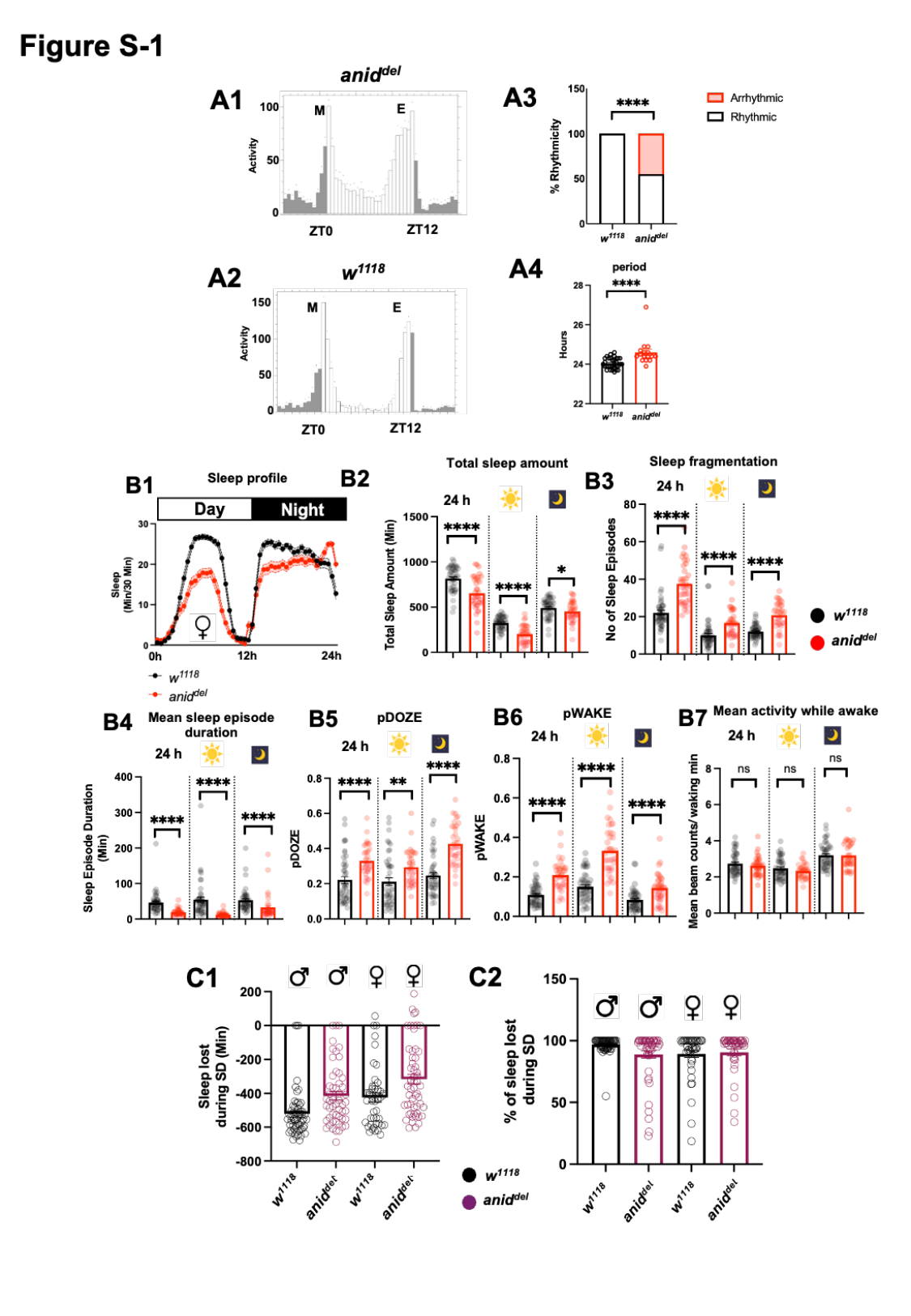

#### Slide 2
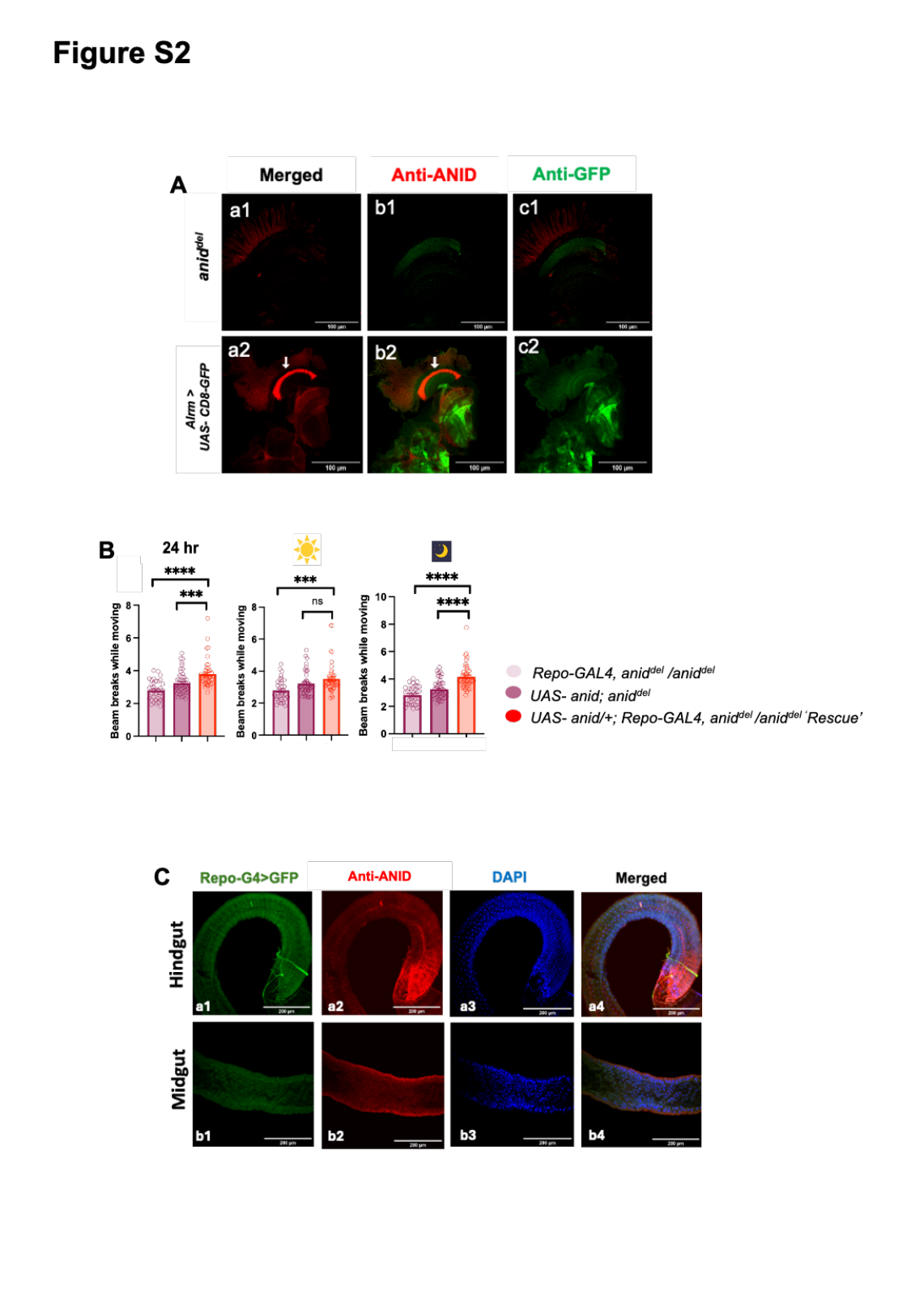

#### Slide 3
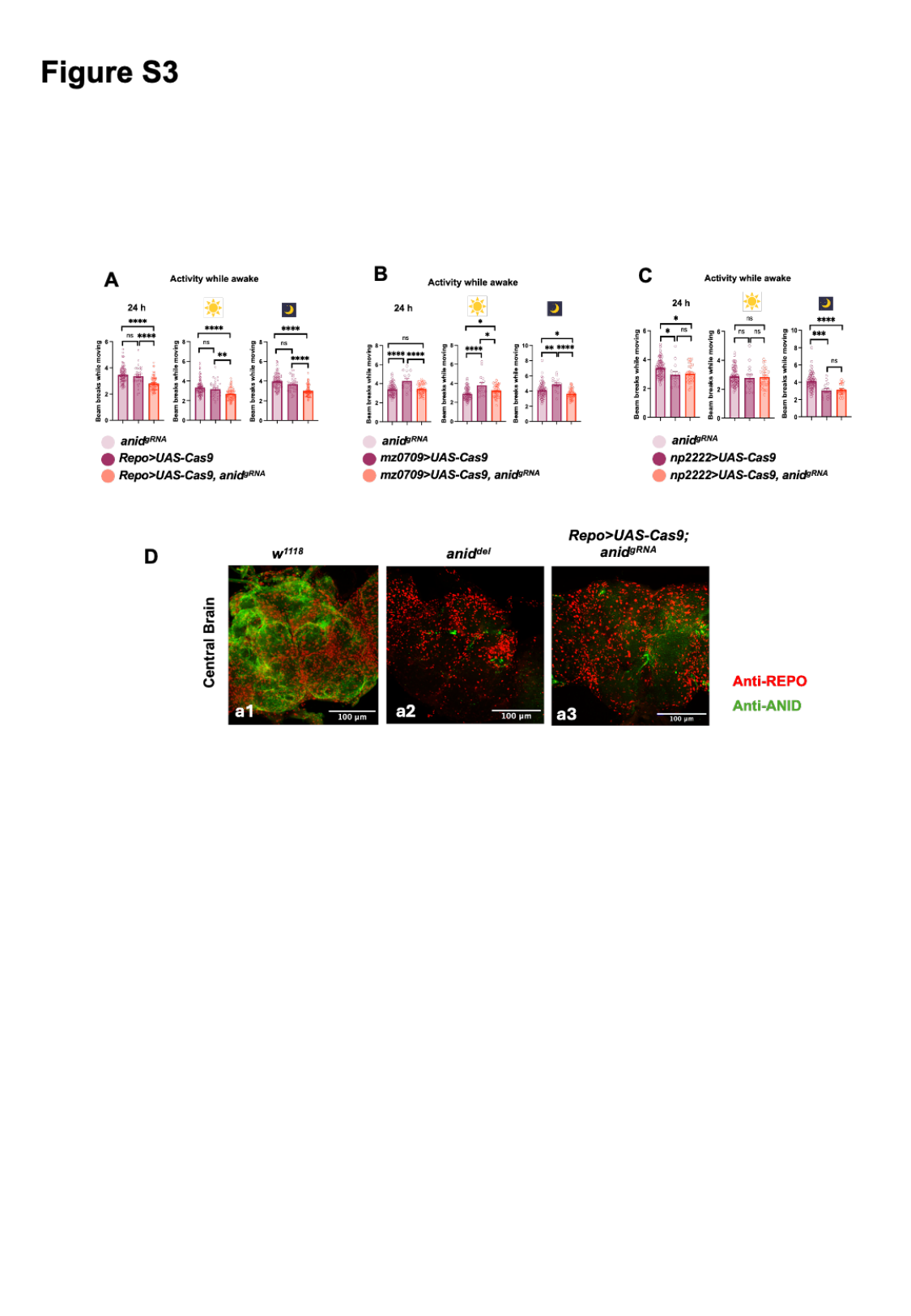

#### Slide 4
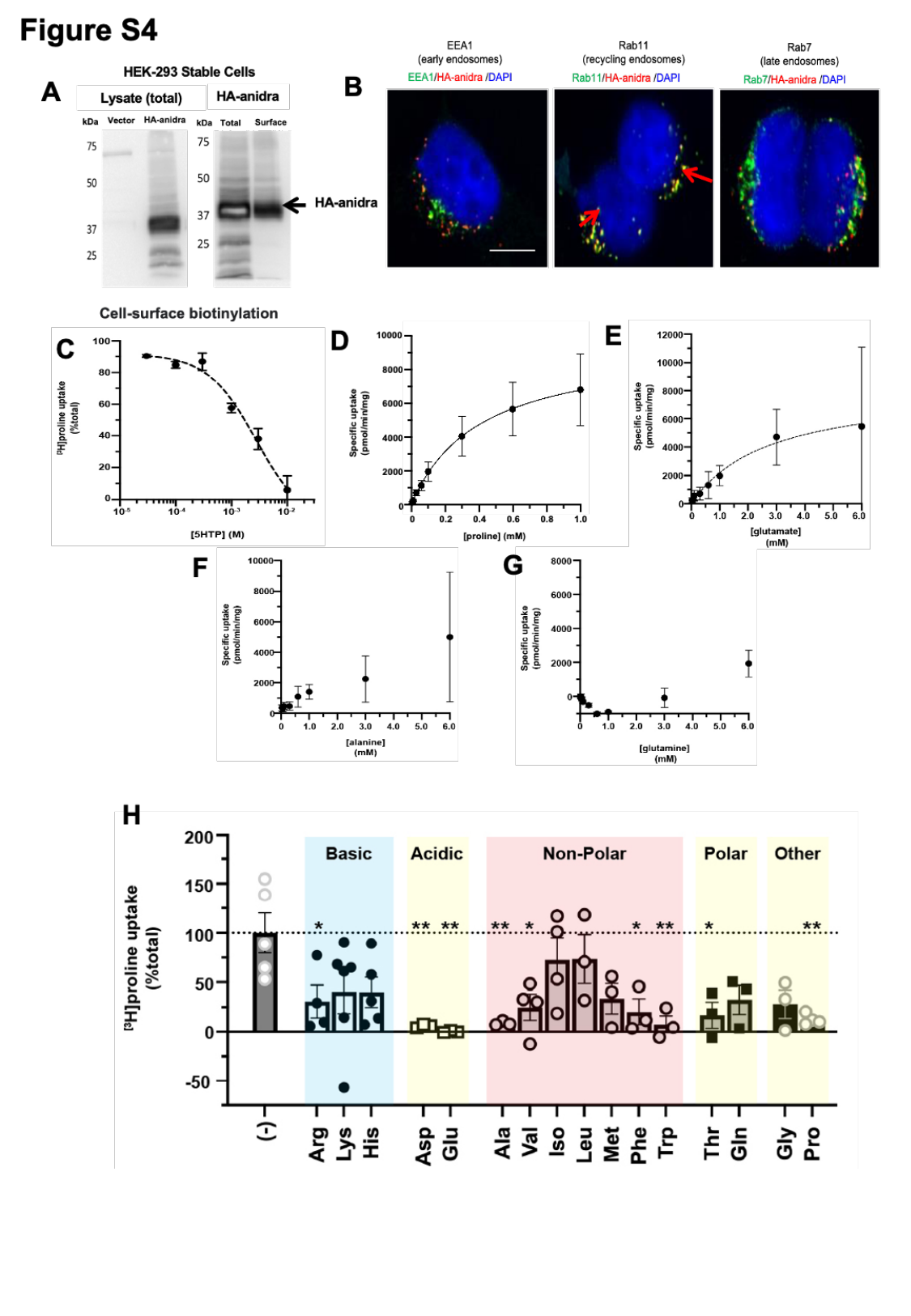

#### Slide 5
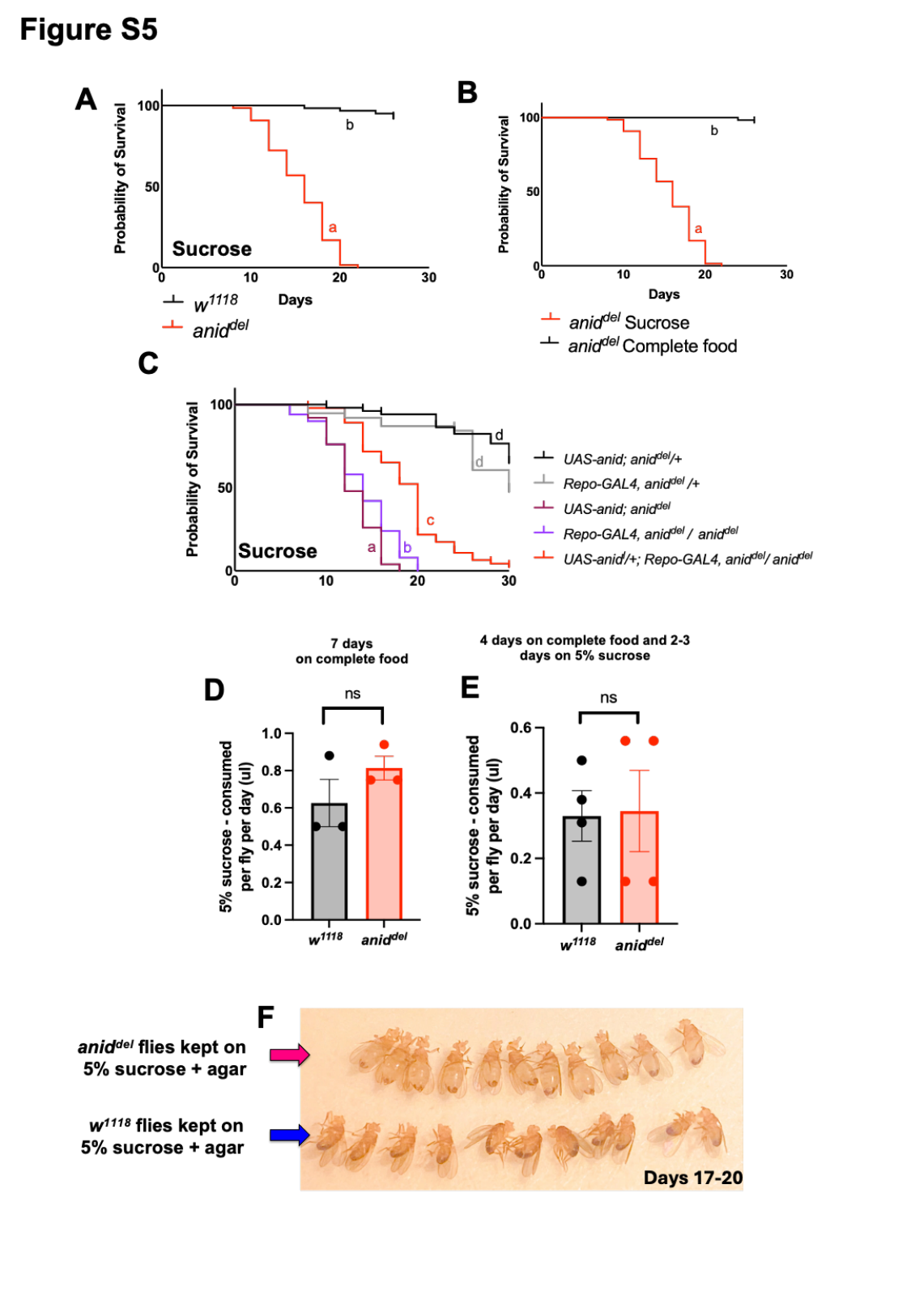

#### Slide 6
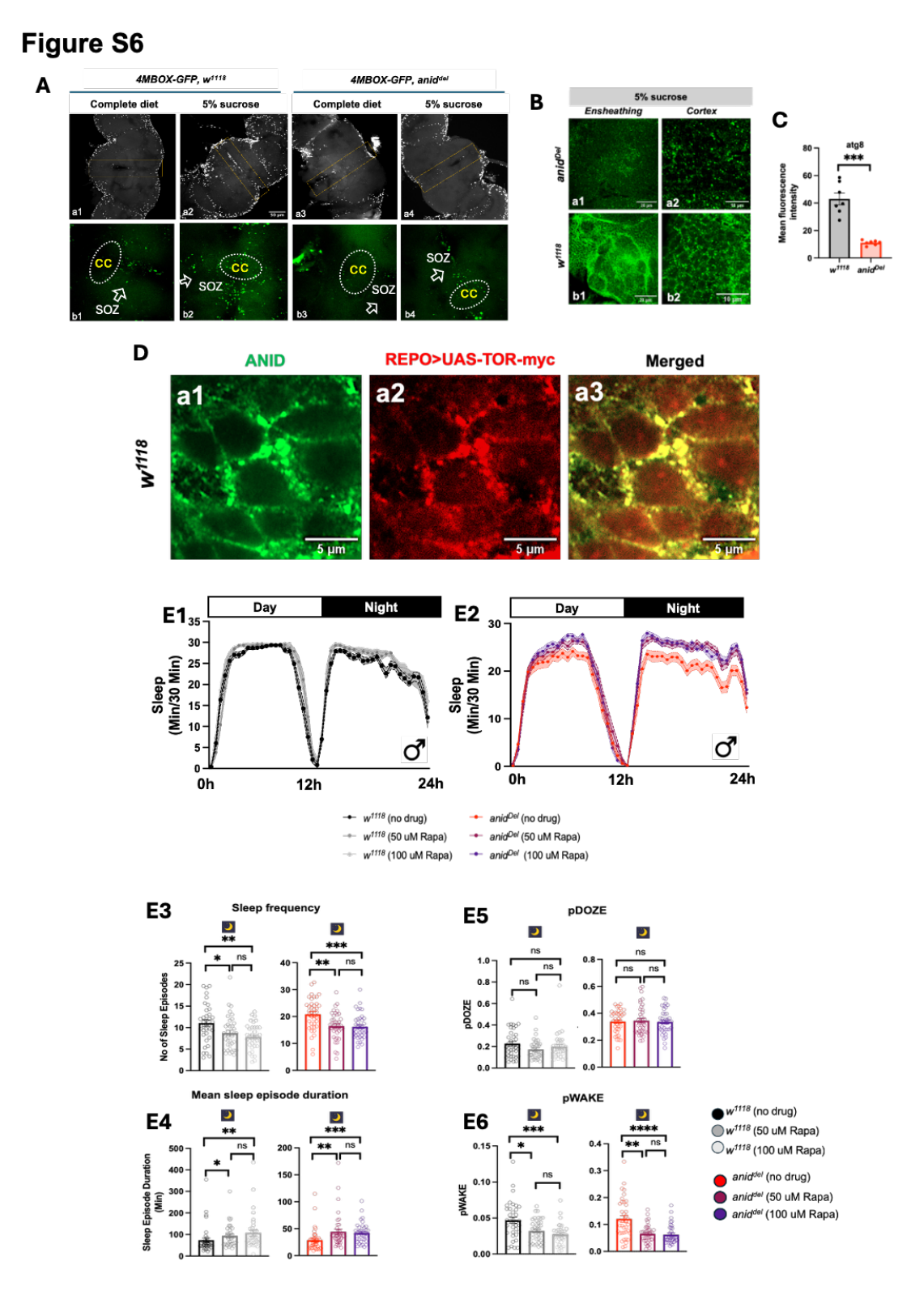
